# DeSide: A unified deep learning approach for cellular decomposition of bulk tumors based on limited scRNA-seq data

**DOI:** 10.1101/2023.05.11.540466

**Authors:** Xin Xiong, Yerong Liu, Dandan Pu, Zhu Yang, Zedong Bi, Liang Tian, Xuefei Li

## Abstract

Cellular decomposition employing bulk RNA-sequencing (RNA-seq) has been consistently under investigation due to its high fidelity, ease of use, and cost-effectiveness compared to single cell RNA-sequencing (scRNA-seq). However, the intricate nature of the tumor microenvironment, and the significant heterogeneity among patients and cells have made it challenging to precisely evaluate the cellular composition of solid tumors using a unified model. In this work, we developed DeSide, a deep learning and single-cell decomposition method for solid tumors, to estimate proportions of cell types presented in tumor samples. Our new deep neural network (DNN) architecture considers only non-cancerous cells during the training process, indirectly calculating the proportion of cancerous cells. This approach avoids directly handling the often more variable heterogeneity of cancerous cells, and instead leverages scRNA-seq data from three different cancer types to empower the DNN model with a robust generalization capability across diverse cancers. Additionally, we used a new sampling method and filtering strategies to simulate the gene expression profiles (GEPs) of solid tumors, creating a carefully controlled training set that could be compared to the bulk RNA-seq data from The Cancer Genome Atlas (TCGA), a database of bulk RNA-seq data collected from cancer patients. Relying on limited yet diverse scRNA-seq data, our approach outperformed current methods in accurately predicting the celluar composition of samples from TCGA and an additional validation set. Furthermore, we demonstrated that the predicted cellular composition can be utilized to stratify cancer patients into different groups with varying overall survival rates. With increased availability of scRNA-seq data for various types of tumors, DeSide holds the potential for a more precise cellular decomposition model using bulk RNA-seq.

## Introduction

Solid tumors consist of various cell types, including not only cancerous cells but also other cell types present in the tumor microenvironment, such as cancer-associated fibroblasts (CAFs), endothelial cells, tumor-associated macrophages, and tumor-infiltrating lymphocytes (TILs) (1–5). In some types of cancer, such as triple-negative breast cancer (TNBC), colon cancer, non-small cell lung cancer (NSCLC), and others, the abundance of TILs has been shown to correlate with the overall survival (OS) or disease-free survival (DFS) of patients (6, 7). As such, accurately evaluating the cellular composition of solid tumors is crucial for understanding the complex tumor microenvironment and its clinical applications, such as patient stratification and identification of therapeutic targets and strategies (3, 6–9).

Currently, the direct evaluation on the cellular composition of solid tumors mainly relies on flow cytometry (10), single-cell RNA sequencing (scRNA-seq) (11), or immunohistochemistry (IHC) analysis of the tumor section images (12). Nevertheless, the cost, coverage of cellular types, or extraction efficiency of cells from solid tumors can all affect the accuracy of evaluation. On the other hand, bulk RNA-seq is widely used for transcriptional level studies of solid tumors at a lower cost. And RNA-seq data of over 10,000 tumor samples is publicly available from the TCGA project (13), which offered a diverse and rich pool of samples for the study of cellular composition of solid tumors, as long as a precise estimation algorithm is available.

In addition, compared to non-cancerous cells within the same tumor microenvirnment, cancerous cells usually experience more disturbance, such as DNA damage, genomic instability, or aberrant DNA methylation patterns (14–16). These genetic and epigenetic changes could contribute extra heterogeneity at gene expression level. As a result, the GEPs of cancerous cells usually exhibit higher variance than those of non-cancerous cells across patients or datasets, as measured in scRNA-seq (**Supplementary Fig. S1**). This wide variation in GEPs across different cancer types makes it impossible to define a universal set of marker genes for all cancerous cells, as we have done for other cell types. Besides, scRNA-seq data for some rare cancer types are insufficient or unavailable, making it much more challenging to represent the heterogeneity of gene expression patterns of diverse cancerous cells across different cancer types compared to non-cancerous cells.

To infer the cellular composition from bulk RNA-seq data, several computational methods have been developed. For one class of deconvolution methods, such as EPIC (17), MuSiC (18), and CIBERSORT / CIBERSORTx (19, 20), a group of characteristic genes is defined for each cell type, and most of them assume that there is a linear relation between the cellular proportion and the representative gene expression profile (GEP) consisting of the characteristic genes. However, these methods have three potential problems: 1) Strong heterogeneity of GEPs within each cell type, particularly in cancer cells, cannot be represented properly by one representative GEP; 2) The optimal gene list for each cell type is hard to determine, and the correlations among these genes are ignored; 3) The wide numerical range of gene expression values in non-log space, usually varies from 0 to tens of thousands of values, and high dimensionality often make the calculation procedures unstable and may produce non-unique solutions, as in the case of non-negative matrix factorization problems (21). To overcome the aforementioned challenges, a decision tree model called Kassandra has been developed (22). However, to achieve higher estimation accuracy, random noise sampled from a Possion distribution needs to be introduced, and the size of the training set required for this method is up to 18 million samples, which can be difficult for regular users to generate and handle.

Alternatively, deep learning algorithms can efficiently capture the intricate structure of large datasets without requiring feature selection (23). At the same time, a pool of scRNA-seq data can potentially preserve the variation of expression patterns within each cell type. Several deep learning-based methods have been developed by combining scRNA-seq data (24–26). These methods use the entire list of measured genes and naturally introduce the cellular heterogeneity. However, by using the scRNA-seq data from homogeneous sources (originating from the same tumor type or the same organ), a simple application of these deep learning-based methods is still unable to accurately estimate the cellular composition of multiple bulk solid tumors by a unified model. This is far from ideal, as we would want to train the model once and use it for all tumor types. Furthermore, it has been claimed that deep learning-based methods may not accurately predict the cellular composition due to the need for a significant training set and the complex and unwieldy training process, which may lead to overfitting (22).

In this work, we present DeSide, a DEep learning and SIngle cell based DEconvolution method, to decipher the cellular composition of solid tumors using low-cost bulk RNA-seq data. To enhance prediction accuracy, we created a comprehensive scRNA-seq dataset by combining publicly available scRNA-seq data from multiple solid tumors and cancer cell lines, and introduced the segment sampling method and data filtering steps to create GEPs of synthetic tumor samples that mimic real tumors with high similarity. We also designed a new DNN architecture using the sigmoid function as the activation function in the output layer to ensure the independence of the predicted cell proportions. With this approach and the high-quality synthetic dataset as a training set, our work offers a method that can accurately estimate the cellular composition of solid tumors using bulk RNA-seq data, even for those tumors whose GEPs may not be available in scRNA-seq data. Comparing the correlations between predicted tumor purity and the consensus measurement of purity estimations (CPEs (27)) across existing methods, we demonstrated that DeSide had the highest concordance correlation coefficients (CCCs) for tumor purity estimation in 14 out of 18 types of cancer in TCGA. DeSide’s accuracy was further confirmed in another recently published bulk RNA-seq dataset of solid tumors with the ground truth of cellular composition verified by flow cytometry. In addition, we also demonstrated that the accurate estimation of cell proportions can help to stratify patients based on their overall suvival (OS). More importantly, our method can be easily expanded to incorporate broader cellular and patient heterogeneity once high-quality scRNA-seq data for other cancer types become available, which could further improve the accuracy of prediction for more tumor types.

## Results

### A new DNN architecture for cell proportion

Considering that the GEPs of cancer cells can be highly variable across individuals and multiple types of solid tumors, while the GEPs of non-cancerous cells are generally more conserved (**Supplementary Fig. S1**), we proposed a new DNN architecture to first predict the proportions of non-cancerous cells **y** within a bulk sample and then deduce the proportion of cancer cells by 1−Σ (y) (**Fig. 1a**). To achieve this goal, instead of using the softmax function (28) which is a common activation function for regression models, we utilized the sigmoid function (28) as the activation function of the output layer. Consequently, the predicted proportion of each cell type is essentially independent of each other (see **Methods**), and the sum of y could be less than one. Using this approach, we first trained the DNN model to accurately predict the proportions of non-cancerous cells. Then, we applied this model to other bulk tumor samples, which may have different tumor types than the ones present in our collected scRNA-seq datasets. As we shall demonstrate in later sections, although the training set was constructed using dataset S1 derived from only three types of cancer, the DNN model with this new architecture enables us to more accurately estimate the tumor purity for multiple tumor types, including those with rare or unavailable corresponding scRNA-seq data.

**Fig. 1.**
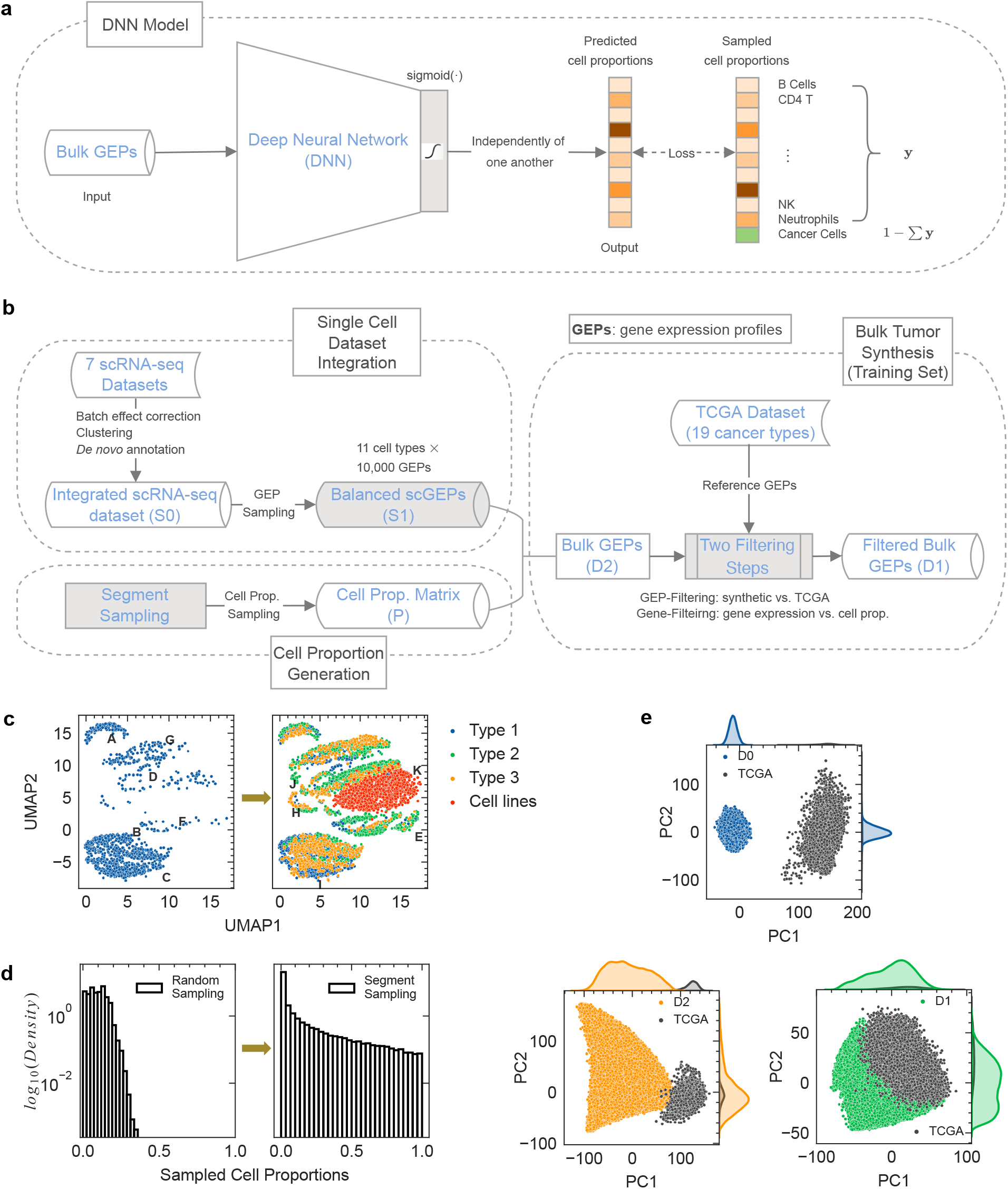
Overview of DeSide. **a**, The DNN model. A fully connected DNN model was used to predict cell proportions of 10 non-malignant cell types (**y**) from bulk GEPs. The cell proportion of cancer cells was calculated by subtracting the sum of **y** from 1. The sigmoid function was used as the activation function of the output layer. **b**, The workflow of bulk tumor synthesis, which includes single cell dataset integration, cell proportion generation, and bulk GEP simulation. The notation of each dataset was marked as Dx or Sx (**Table S1**), and the key steps were highlighted with a grey background. The integrated scRNA-seq dataset, S0, contains 11 re-annotated cell types after batch effect correction and clustering. S1, which was generated from S0, and the cell proportion matrix (P) were necessary to generate bulk GEPs. The TCGA dataset was used as a reference dataset to evaluate the quality of generated bulk GEPs. **c**, UMAP visualization of the integrated scRNA-seq dataset S0. 6 out of 7 scRNA-seq datasets originated from 3 different solid tumor types (Type 1 - Type 3), and each dataset contains multiple cell types. The other dataset is a pan-cancer scRNA-seq dataset, which contains multiple cancer cell lines from 22 different cancer types. The re-annotated cell types were marked as capital letters A-K. **d**, The distribution of generated cell proportions of a single cell type (other cell types have similar distributions) by different sampling methods. Left: random sampling from a uniform distribution in the interval [0,1), which can only generate low proportions for all cell types. Right: segment sampling can generate up to 100% proportion for any cell type. **e**, The PCA visualization of simulated datasets (training sets) and the TCGA dataset. For each panel, one of the three simulated datasets D0, D1, or D2 was combined with TCGA to perform PCA analysis.

### Novel strategies to synthesize GEPs of solid tumors

To prepare high-quality training sets, we used new strategies to synthesize GEPs that mimic the bulk RNA-seq data of solid tumors as closely as possible, as summarized in the workflow shown in **Fig. 1b**.

More specifically, despite recent advances and broad applications of scRNA-seq, an individual dataset collected from cancer patients usually includes only a few cell types, and the number of cells for certain types also could be limited (**Fig. 1c**, left panel). Furthermore, compared to bulk RNA-seq data, scRNA-seq data are generally more noisy and variable. To overcome these challenges, we created a comprehensive dataset (S1) with 11 cell types and 10,000 single-cell gene expression profiles (scGEPs) for each cell type, where each scGEP is essentially the average gene expression profile of 100 single cells with the same type (see **Methods**). In addition, UMAP visualization of dataset S1 showed that it captured most of the gene-expression diversity of each cell type (**Supplementary Fig. S12**). Therefore, a balanced dataset of scGEPs was created, which were used to computationally synthesize bulk tumor samples in the following steps.

In addition to creating the comprehensive dataset S1, we also proposed a new sampling method for generating cell proportions, as well as two filtering steps for synthesized bulk GEPs. We will provide a brief illustration of the procedures and effects of the proposed strategies in this section.

Specifically, we first generated a cell proportion matrix (**P**, also see **Fig. 1b**) to define the proportion of each cell type in each synthetic tumor. For this step, most of previous methods used a sequential random sampling method (24–26) from a uniform distribution. However, we found that this sampling method can only generate cell proportions with low proportions for all cell types (**Fig. 1d**, left). In this work, we used a new sampling strategy, segment sampling, to generate the cell proportion matrix. And we demonstrated that the distribution of the percentages of each cell type can cover the whole range from 0 to 1 (**Fig. 1d**, right panel, see **Methods**).

Furthermore, to increase the similarity between bulk GEPs of training set (GEPs of synthetic tumors) and bulk RNA-seq data from TCGA dataset for a better training performance, we introduced “GEP-level filtering” and “gene-level filtering” for the synthesized bulk GEPs (see **Methods** and **Fig. 1b**).

To compare the diversity and quality of generated datasets by different methods and demonstrate the necessity of the new sampling strategy and filtering steps, we compared these datasets with the TCGA dataset by PCA analysis (**Fig. 1e**). In the PC1-PC2 space, the dataset generated using random sampling without filtering (D0) was found to be separated from TCGA, while the synthetic GEPs generated using segment sampling without filtering (D2) had a small overlap with TCGA and left most of the area of TCGA uncovered. For the effects of filtering steps, the synthetic dataset generated using segment sampling with filtering (D1) showed a large overlap with and covered most of the area of TCGA. To track the effect of different datasets further, we input each training set and TCGA data into the corresponding DNN model after training, and used the output (GEP embedding, a low-dimensional vector) of the first hidden layer to perform another PCA analysis (Supplementary **Fig. S2** and Methods). The PCA visualization revealed that the GEP embedding of D1 in the first hidden layer had an even better coverage for the space taken by that of the TCGA data, while D0 had little overlap with TCGA and D2 had a mismatch with TCGA in the high-density area.

These results collectively suggested that the improved sampling method enables the synthetic bulk GEPs to have a broader coverage, and the filtering steps further increase the similarity of gene expression profiles between synthetic tumors and real tumor samples from TCGA. The detailed comparison of the training efficiency among various training sets will be further discussed in later sections.

### The GEP diversity of training set can affect the coverage of DNN models

When using machine learning models to decompose the cellular proportions in bulk samples, one of the challenges is that the GEPs in training sets can be different from the bulk RNA-seq data of real samples (22, 26). To prepare an appropriate training set, we compared the performance of the DNN models that were trained by various training datasets in **Fig. 2a**. Note that all DNN models shared the same architecture. In addition to dataset D0, D1 and D2, we also created a training set (D1+D2) by combining D1 and D2.

**Fig. 2.**
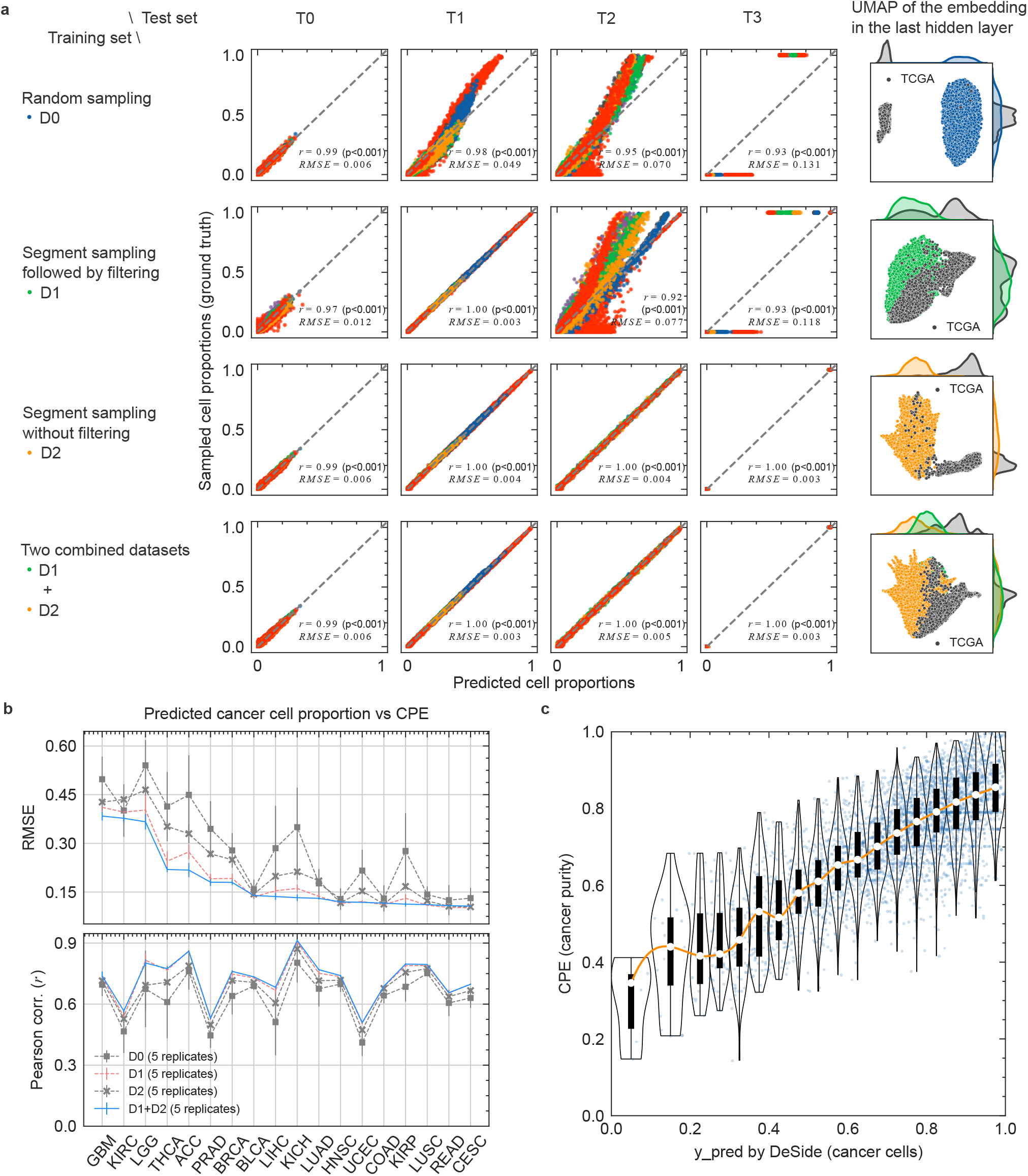
DNN models exhibit varying performance when trained on different training sets. **a**, Four DNN models were evaluated on four different test sets (T0 to T4). All models have the same architecture but were trained on different datasets generated by various methods: random sampling (D0), segment sampling followed by filtering (D1), segment sampling without filtering (D2), and a combination of D1 and D2 (D1+D2). Different colors in the 16 panels represent different cell types. The similarity between the training set and TCGA in the last hidden layer of the corresponding DNN model is visualized in the last column by UMAP. **b**, The root mean square error (RMSE) and Pearson correlation between predicted cancer cell proportions and cancer purity (CPE value (27)) were calculated for 18 tumor types (PAAD has no CPE). **c**, Comparison of the DeSide-predicted cancer cell proportions with the CPE values for 10 tumor types using analysis criteria of *r ≥* 0.6 and *RMSE ≤* 0.15, as shown in (b). In the violin plots, the data were manually binned according to the predicted cancer cell proportions on the x-axis, and each sample with a CPE value was denoted by a light blue dot. The upper and lower edges of the black bars represent the upper and lower quartiles of the data, respectively, while the white circles indicate the median CPE values in each bin. The orange line is used to connect the median values and is smoothed.

In total, we used 4 different training sets (D0, D1, D2 and D1+D2, **Fig. 2a**) and tested the performance of each DNN model on 4 different test sets (T0 - T3, **Fig. 2a**). For the test sets, specifically, test set Ti was generated by the same method as the training set Di, where *i ϵ* {0, 1, 2}. Test set T3 consists of GEPs with single cell type (SCT), which were generated by the same method as dataset S1 (scGEPs, see **Methods** and **Table S1**). We used T3 to test an extreme situation in GEPs, where one cell type had 100% cell proportions and all other cell types were not involved.

Not surprisingly, the training sets strongly affected the performance of the DNN models. As shown in **Fig. 2a**, the model always performed well (high correlation and low RMSE error) on the test set that was generated by the same method used for generating the corresponding training set (the first 3 subplots on the diagonal of **Fig. 2a**). The DNN model trained by D0 or D1 showed low performance on the test sets that were generated by different methods. Interestingly, the DNN model trained by D2 showed excellent performance across all 4 test sets, which suggested that the method to generate synthetic tumors in D2, i.e. segment sampling without filtering, could cover a broader GEP space compared to methods that generated D0 and D1.

### The GEP similarity between training set and real tumors can strongly influence the performance of DNN models

Although the training set D2 showed broad diversity and the DNN model trained by D2 performed well for all 4 test sets, we will show in the following section that when estimating the cellular composition of real tumors, the similarity between synthetic GEPs and real bulk RNA-seq data also plays a significant role.

To evaluate the similarity between synthetic bulk GEPs and bulk RNA-seq data from TCGA, we first compared the UMAP visualization of various training sets and the TCGA dataset in the last hidden layer of each DNN model, as shown in the last column of **Fig. 2a** (similar to Supplementary **Fig. S2**, also see **Methods**). As expected, for DNN models that included D1 as a part of the training set (row 2 and row 4 in **Fig. 2a**), where the synthetic GEPs were filtered to be similar to bulk RNA-seq data in TCGA (see **Methods**), the area occupied by TCGA data was mostly covered by that of synthetic GEPs.

Furthermore, to demonstrate the effects of various training sets on the estimation accuracy of the cellular composition of real tumors, we performed the Pearson correlation analysis between the cancer cell proportions predicted by the aforementioned 4 DNN models and CPE values for 18 tumor types (see **Methods**). As demonstrated in **Fig. 2b**, the DNN model trained by the combined D1 and D2 dataset (D1+D2) consistently showed the highest correlation and lowest RMSE error for most of tumor types studied.

In addition, since there is no “ground truth” in the bulk RNA-seq data from TCGA for the DeSide-predicted proportions of non-cancerous cells in real tumor samples, we employed a signature score based on the average expression values of marker genes (see **Methods**) to indicate the relative abundance of non-cancerous cells. We further showed the Pearson correlation coefficients between the predicted cell proportions by the aforementioned 4 DNN models and the corresponding signature scores based on the bulk RNA-seq data for each cell type across all 19 cancer types. Interestingly, DNN models trained by datasets that included D1 had significantly higher correlation coefficients compared to those in which D1 was not included as part of the training set (Supplementary **Fig. S3**). For most cell types, the model trained by the dataset D1+D2 has similar or higher correlation than other models.

Therefore, we would like to argue that the similarity between synthetic GEPs and bulk RNA-seq data in TCGA is essential for accurately estimating the cellular proportions. The results for all cancer types that predicted by DeSide using D1+D2 as training set can be found in the first column of Supplementary **Fig. S4** - **S5**. And in all following sections, we will demonstrate the performance of DeSide using the D1+D2 dataset.

### Improved cellular composition estimation using DeSide and CPE values with calibration

Although De-Side has not yet achieved the perfect accuracy in predicting tumor purity, for an unseen bulk sample, it can provide a reasonably good estimation of its cellular composition by considering the CPE values of TCGA bulk samples. Specifically, for each TCGA tumor sample (7,699 in total) considered in this study, DeSide predicted the tumor purity, as well as the CPE value was estimated in another study (27). By plotting the two sets of values together (**Fig. 2c**, light blue dots in the background), we generated the probability distribution of CPE values within each section of DeSide-predicted tumor purity (**Fig. 2c**, violin plots). The width of each section was adjusted manually, taking into account the number of available samples in each section. Consequently, given the DeSide-predicted tumor purity of an unseen bulk sample with bulk RNA-seq data, its most-likely CPE value can be obtained within the 95% confidence interval (CI) (**Table S6**). Moreover, after properly estimating the tumor purity, the absolute proportions of non-cancerous cells can also be corrected based on the ratios between non-cancerous cells estimated by DeSide. These calibrations allow for a more accurate estimation of the cellular composition in real samples using DeSide and CPE values of TCGA tumor samples.

### Performance comparison between DeSide and other algorithms

To compare the performance between DeSide and other algorithms, we first predicted the cancer cell proportions of 18 tumor types in TCGA by Scaden, EPIC, CIBERSORTx, DeSide, and Kassandra (see **Methods**). The concordance correlation coefficient (CCC) between predicted cancer cell proportions and CPE values was calculated to evaluate the performance across different algorithms (**Fig. 3a**). The Pearson correlation (*r*) and root-mean-square error (RMSE) are also shown in Supplementary **Fig. S4** -**S5**. The results demonstrate that DeSide achieved the highest performance for 14 out of the 18 tumor types studied in this work. As expected, DeSide consistently outperformed methods that use signature genes, such as CIBERSORTx and EPIC. After removing two negative controls, i.e., GBM and LGG, we compared the difference of CCCs between DeSide and all other algorithms by Wilcoxon signed-rank test (**Fig. 3b**). DeSide outperformed all these algorithms significantly.

**Fig. 3.**
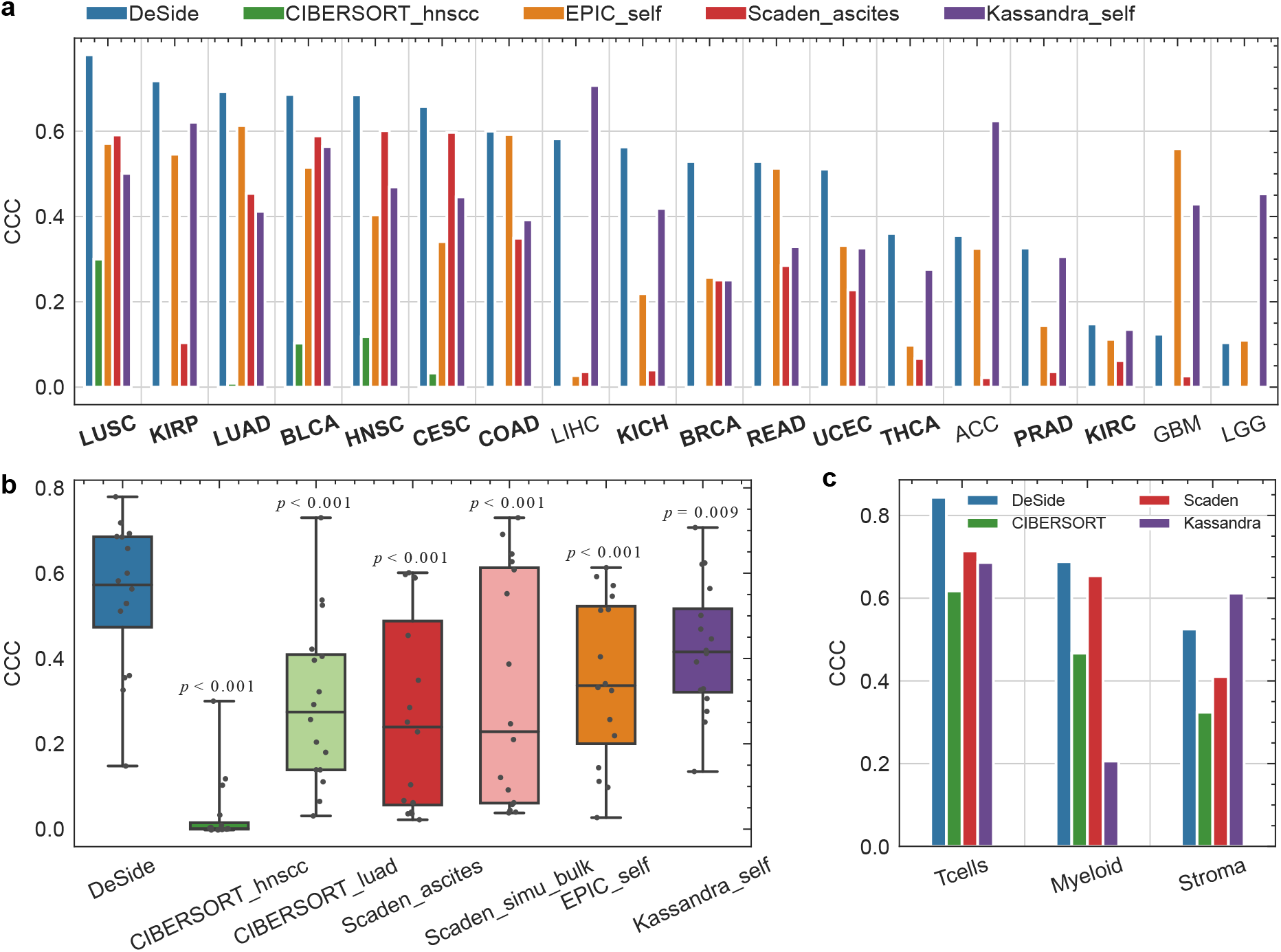
Performance comparison between DeSide and other algorithms. **a**, Comparison of concordance correlation coefficients (CCCs) between predicted cancer cell proportions and cancer purity (CPE values) (27) across 18 tumor types using different algorithms. The cancer types where DeSide performed the best are highlighted in bold, and the list of cancer types was sorted by the CCCs of DeSide in descending order. **b**, Comparison of CCCs among different algorithms or datasets in 16 cancer types after removing two negative controls, i.e., GBM and LGG. The p-value for each comparison was calculated using Wilcoxon signed-rank test with DeSide as the reference. **c**, Comparison of CCCs between predicted cell proportions and flow scores on solid tumor samples in dataset GSE184398 (29) across three different cell types: Tcells (CD4^+^ and CD8^+^ T cells), Myeloid (DC, macrophages, neutrophils, and mast cells), and Stroma (fibroblasts and endothelial cells). Both the predicted cell proportions and flow scores for each cell type were normalized by dividing the corresponding maximum values (see Methods). The suffix “_simu_bulk” indicates that the dataset D1+D2 was used as the training set. The suffix “_ascites” indicates that the ascites dataset provided by Scaden was used as the training set. The suffixes “_hnscc” or “_luad” indicate that the scRNA-seq data originated from HNSCC (provided by CIBERSORT) or LUAD (provided by this study) was used as the reference dataset to generate the signature matrix. Further details on the algorithm comparison can be found in the Methods section.

In addition, we compared the performance of CIBER-SORTx and Scaden using different datasets (**Fig. 3b**, Supplementary **Fig. S6**). When using the scRNA-seq dataset LUAD (provided by this study) as the reference dataset to generate the signature matrix for CIBERSORTx, we observed a significant improvement in performance for all cancer types compared to the original signature matrix generated using the scRNA-seq dataset HNSCC (provided by CIBERSORTx) (light and dark green bars in Supplementary **Fig. S6**), especially for the cancer type LUAD, which has the same type as the provided scRNA-seq dataset. For Scaden which also employed deep learning and scRNA-seq data, we observed a similar situation. When we used the dataset simu_bulk (i.e., D1+D2) as the training set, the performance of Scaden improved for most cancer types compared to using the dataset ascites (provided by Scaden) as the training set (light and dark red bars in Supplementary **Fig. S6**). Interestingly, even using the same training set (simu_bulk), the accuracy of Scaden was worse than that of DeSide for most cancer types (Supplementary **Fig. S6**). This suggests that the architecture of the DNN model used by DeSide may help to increase the estimation accuracy as well. We also demonstrated that using sigmoid function as the activation function of the output layer and the “1 − Σ (**y**)” strategy improved the performance of DeSide among most of cancer types further (Supplementary **Fig. S6**). For the most recent method, Kassandra, considering the size of its training set (18 million for Kassandra-Tumor model) compared to DeSide (0.2 million), the superior efficiency of DeSide in learning valid information from the training set is apparent.

The above comparison indicates that finding a suitable dataset for a specific task can be crucial and challenging. Therefore, a better strategy could be to develop a unified model that considers as many similar cancer types as possible as a whole. On the other hand, we would like to emphasize the generalisation ability of DeSide that was trained by the combined dataset D1+D2. For LUAD and HNSCC, the scRNA-seq data were used to generate the training set, and therefore we would expect a reasonably good correlation between the DeSide-predicted tumor purity and the CPE values. Indeed, we observed that the correlation coefficients for LUAD and HNSCC were above 0.6 and the RMSEs were less than 0.15 (Supplementary **Fig. S4** -**S5**). Including these two, 10 out of 18 tumor types studied in this work showed a good correlation (≥ 0.6) and low RMSE (≤ 0.15) between the DeSide-predicted tumor purity and the CPE values. Furthermore, for GBM and LGG, which are both brain tumors with the neuroepithelium origin and are used as negative controls since they differ from other cancer types originating from epithelial cells, DeSide-predicted tumor purity also showed a high correlation with CPE values, although with a higher RMSE (*>* 0.35). Interestingly, we noticed a systematic shift between the predicted tumor purity and CPE values, which could be attributed to the distinctive brain tumor microenvironment characterized by its unique tissue-resident cell types (30) and the presence of the blood-brain barrier (31).

Therefore, DeSide demonstrated fairly good generalisation ability and indeed learned some common structures shared by multiple solid tumors, which is notable considering that the collected scRNA-seq dataset used for the training set contained multiple different tumor types, i.e., PAAD (without CPE values), LUAD and HNSCC. Although the diversity and sample size of this dataset are still limited, this allowed the model to learn the commonalities across multiple tumor types and improve its ability to generalize to new data.

To further demonstrate the predictive power of DeSide on the cellular proportion of non-malignant cells in bulk samples, we applied DeSide to another recently published dataset, GSE184398, with the “ground truth” of cellular proportions measured by flow cytometry (29). This dataset contains 12 different solid tumor types obtained from 364 fresh surgical specimens. 260 bulk RNA-seq profiles were downloaded from the GEO database (GEO184398-Live, i.e., all viable cells at the time of sorting), and for each sample, the corresponding flow scores were determined by flow cytometry for a few types of cells. DeSide showed the highest CCC values for T cells and myeloid cells, and the second highest CCC value for stromal cells (**Fig. 3c**). The CCC value, Pearson correlation, and RMSE between predicted proportions and the flow scores of each cell type were shown in Supplementary **Fig. S7** (see **Methods**).

### Patient stratification through DeSide-predicted cellular proportions

It has been demonstrated that the abundance of specific types of cells in solid tumors can help stratify patients into distinct prognostic groups with different overall survival (OS) and/or disease-free survival (DFS) rate (32, 33). To investigate the efficacy of DeSide in stratifying patients, we tested whether the cell proportions estimated by DeSide can separate patients into significantly different prognostic groups for 10 types of tumors (COAD, KIRP, CESC, BLCA, LIHC, HNSC, KICH, LUAD, LUSC, READ). These 10 tumor types were selected based on the criteria that the Pearson correlation coefficients (*r*) between predicted tumor purity and CPE values were higher than 0.6, and the RMSEs were lower than 0.15.

First, by considering the proportion of only one cell type, we separated patients with the same cancer into two groups using the median value of predicted cell proportions. For a few types of tumors, our results demonstrated that the proportions of fibroblasts, CD8^+^ T cells, CD4^+^ T cells, macrophages, or B cells can indeed stratify the patients into two groups with significantly different Kaplan-Meier overall survival (OS) curves (**Fig. 4a** and Supplementary **Fig. S8a**). For example, HNSCC and BLCA patients with a higher proportion of B cells and lower proportion of fibroblasts have a longer OS, respectively. Consistently, similar trends were observed in recent studies (32, 34, 35).

**Fig. 4.**
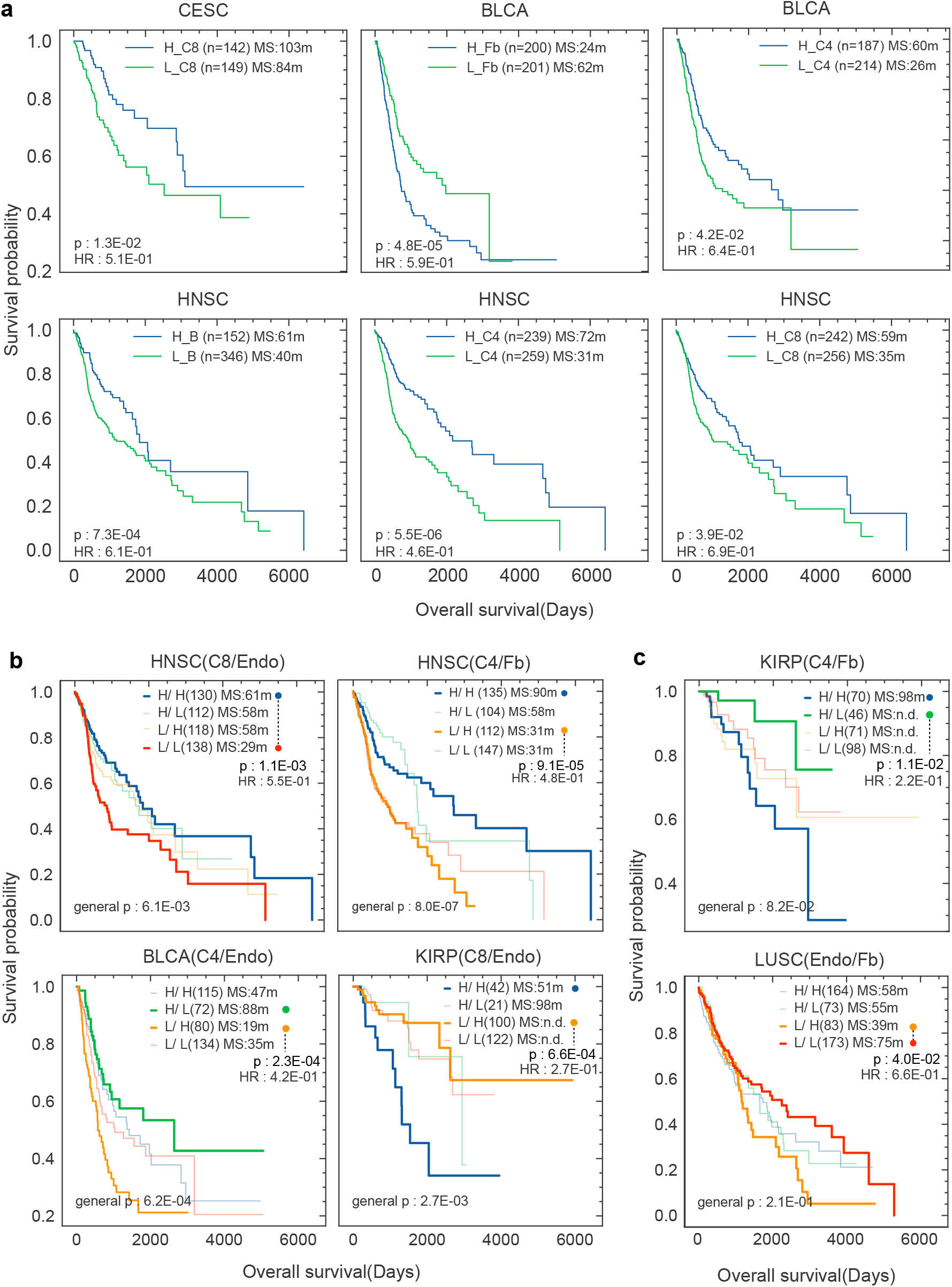
Kaplan-Meier overall survival (OS) curves for TCGA cohort stratified by the cell proportions predicted by DeSide. **a**, OS curves showed significant differences when classified by the cell proportions of a single cell type. The p-values were calculated by the log-rank test. **b**, OS curves showed significant differences when classified by the cell proportions of two cell types. Situations where one of the cell types in each pair could also be able to separate patients into two groups are shown in (a) or (Supplementary **Fig. S8a**). **c**, OS curves showed significant differences when classified by the cell proportions of two cell types, where both cell types were not shown in (a) or (Supplementary **Fig. S8a**). The bottom left p-values were used to test the overall difference of four survival curves, and the top right p-values were used to test the difference between the best and the worst survival curves. The log-rank test was used for all statistical significance testing. The numbers in parentheses denote sample numbers. MS, Median survival time, measured in months (m); n.d., not determined; HR, Hazard Ratio; B, B cells; Endo, Endothelial Cells; Fb, Fibroblasts; C4, CD4^+^ T cells; C8, CD8^+^ T cells; B, B Cells; H, High; L, Low.

Next, we considered two cell types together and tried to explore the effect of their interaction in stratifying patients. Specifically, for patients with the same type of cancer, we separated them into four sub-groups based on the median cell-proportion values of two cell types (see **Methods**). Then we focused on the best- and worst-survival groups, and tested whether there was any significant difference among them in patient OS (*p <* 0.05, **Fig. 4b, 4c** and Supplementary **Fig. S8b**).

On the one hand, it is not surprising to identify a significant difference between two cell types (**Fig. 4b** or Supplementary **Fig. S8b**), where one of them alone can already stratify patients effectively (**Fig. 4a** or Supplementary **Fig. S8a**). For example, as demonstrated in **Fig. 4b** (upper right), when the patients have higher levels of fibroblasts, the CD4^+^-high group showed the longest survival while the CD4^+^-low group showed the worst prognosis in overall survival among all four groups. Above result is not surprising because the abundance of CD4^+^ T cells alone is already able to stratify patients with significant difference as shown in **Fig. 4a** (bottom middle panel). Moreover, in this specific case, a larger survival difference was observed between the two groups (**Fig. 4b** upper right panel, H-H versus L-H) compared to the survival difference between the two groups stratified by only one cell type (**Fig. 4a** bottom middle panel, H versus L). The difference in median survival time (MS) between these two situations is 59 months for the comparison we discussed earlier (**Fig. 4b** upper right panel, H-H versus L-H) and 41 months for the other comparison (**Fig. 4a** bottom middle panel, H versus L). For the former comparison (**Fig. 4b** upper right panel, H-H versus L-H), the abundance of fibroblasts does not explain the difference in survival between the two groups, as both groups have high levels of fibroblasts. It suggests that the interaction between the two cell types could be one of the factors contributing to the increased gap in survival differences from 41 months to 59 months. Such analyses can help us further screen patients with high risk.

Furthermore, for certain cell combinations in KIRP and LUSC, we found that neither of two cell types’ proportions alone could stratify patients into long- or short-survival groups with significant differences. However, a particular combination of these two cell types could (**Fig. 4c**). For example, under the condition that the patients had a low proportion of endothelial cells, LUSC patients with a lower proportion of fibroblasts survived significantly longer than those with a higher fibroblast proportion (bottom panel of **Fig. 4c**). This finding indicates an interaction between these two cell types, as we previously analyzed. We believe that conducting such analyses, which rely on accurate prediction of cellular composition, can help identify clinically relevant combinations of various cell types and potentially enhance prognostic analyses.

## Discussion

Cellular decomposition of solid tumors with bulk RNA-seq data can offer a quantitative understanding of the tumor microenvironment and tumor heterogeneity in a low-cost fashion. More importantly, such information can help to stratify patients in terms of the prognosis or responsiveness to therapies. So far, there have been a series of efforts to develop an accurate decomposition method using signature-genes or machine learning-based strategies. Although other strategies by combining deep learning and scRNA-seq data existed (24– 26), the current accuracy of such methods is not comparable to the best machine learning-based method, and there is concern about the over-fitting, the size of the training set, and optimization methods (22).

In this work, we properly prepared the training dataset by using publicly available scRNA-seq data from more than 130,000 cells obtained from 105 patients with HNSC, PDAC, or LUAD (see **Table S2-S3**). To the best of our knowledge, this is the first scRNA-seq dataset that combines multiple cancer types and is used to tackle cellular decomposition of solid tumors. Using this dataset and a new architecture, we developed a deep learning-based algorithm called DeSide that can accurately estimate the absolute proportions of 11 cell types in bulk samples. When tested against the tumor purity of TCGA tumors as well as bulk samples with the ground truth of cellular proportions, DeSide demonstrated excellent accuracy compared to currently available methods. Interestingly, we achieved such accuracy only with 200,000 training samples, which is well manageable under the machine learning-based method. This suggests that segment sampling, as well as the GEP and gene-level filtering proposed in this work (see **Methods**), significantly increase the efficiency of the deep learning model.

To better characterise the tumor microenvironment, it is important to not only predict cell proportions in bulk samples but also dissect the gene expression profiles of each cell type. Currently, CIBERSORTx (20) is capable of achieving such deconvolution, however, the estimation based on the same algorithm presented by CIBERSORT (19), which was not accurate enough for a wide range of cancer types, as demonstrated in this study. Additionally, it can only be used for samples with significant differences in GEPs between two groups. With more accurate predictions of cellular composition, DeSide can be a valuable tool for gene expression profile deconvolution when combined with CIBERSORTx or another DNN model. It is even possible to perform both steps, cell proportion estimation and gene expression profile decomposition, using a single DNN model, which could exploit the correlation information between genes.

Furthermore, it has been demonstrated that the spatial distribution patterns of non-cancerous cells are essential for predicting the patient prognosis as well as the responsiveness to the therapies (36). For example, more CD8^+^ T cells in solid tumors does not necessarily mean a good prognosis of patients with triple negative breast cancer, where the infiltration pattern of CD8^+^ T cells into the tumor-cell clusters has to be considered (37). Therefore, the combination of accurately predicted cellular composition by DeSide, precise deconvolution of gene expression profiles of various types of cells, and quantitative characterization of spatial profiles of various types of cells holds promise for more precise patient stratification. More importantly, this approach offers a more systematic picture of the tumor microenvironment from the perspective of cell-cell interactions.

Although DeSide has not achieved perfect accuracy yet, we would like to point out that the procedure used by DeSide is straightforward and can be easily improved and expanded in the future with more scRNA-seq data. In **Fig. 1e**, the gene expression profiles of synthetic tumors still do not cover the entire range of TCGA data, which may reduce the accuracy of DeSide for certain cancer types. One way to overcome such barrier is to include scRNA-seq data from more patients and different cancer types to increase the diversity in the gene-expression space of synthetic tumors. Secondly, DeSide can currently predict the proportions of 11 types of cells. However, scRNA-seq data inherently provides the high resolution of cell types, allowing for additional types or subtypes of cells to be added as long as there are enough cells and the training process is manageable. Thirdly, blood samples have a very different composition compared to solid tumor samples, and the gene expression patterns of immune cells might be distinct from those in the tumor microenvironment. To improve the performance of DeSide on blood samples, transfer learning (38) could be applied.

Nonetheless, our study has shown that DeSide outperforms other methods in accurately predicting the cellular composition of tumor samples across multiple cancer types using bulk RNA-seq data. As more scRNA-seq data becomes available, sampling filtering strategies continue to improve, DeSide has the potential to provide even more precise estimation of cellular composition in a variety of bulk samples.

## Methods

### Information about the collected datasets

We collected two types of datasets: single cell RNA-seq (scRNA-seq) datasets and bulk RNA-seq datasets. The information about the collected scRNA-seq datasets is summarized in **Table S2**. And the cell types included in scRNA-seq datasets are B Cells, CD4 T (CD4^+^ T cells), CD8 T (CD8^+^ T cells), Cancer Cells, DC (dendritic cells), Endothelial Cells, Fibroblasts, Macrophages, Mast Cells, NK (nature killer cells), and Neutrophils. The TCGA dataset is one part of the bulk cell RNA-seq datasets used in our study. And we also collected another bulk RNA-seq dataset (29) with experimental results, which was used as external test set to evaluate the performance of DeSide.

### Integrating single cell RNA-seq datasets

To better represent the cellular heterogeneity across different solid tumors, we created a comprehensive dataset S0 (see **Table S1**) by pooling 7 different scRNA-seq datasets (**Table S2**). 6 datasets were collected from 3 tumor types (head and neck cancer (39, 40), pancreatic ductal cancer (41, 42) and lung cancer (43, 44)). We only used the samples originated from primary tumors and samples came from metastatic tumors or peripheral blood mononuclear cells (PBMCs) were removed. To capture a wider range of cancer cell GEPs, we included another pan-cancer scRNA-seq dataset, which contains 198 cancer cell lines from 22 different cancer types (45). We performed *de novo* cell type annotation for each cell and batch-effect correction for these 7 datasets. The marker genes used for the annotation of cell types were collected from multiple publications (**Table S4**). Following the procedures described in *Scanpy* tutorials (https://scanpy-tutorials.readthedocs.io/en/latest/pbmc3k.html), cells with low quality and cells that belonged to small clusters during the annotation process were removed from the combined dataset. In total, we collected GEPs of 135,049 cells, covering 11 cell types (Supplementary **Fig. S9** - **S11**, **Table S3**).

### The generation of dataset S1

It is well-known that the measurements of scRNA-seq can be highly variable from cell to cell and suffer from sparsity, where the ratios of 0s in one GEP can be as high as 95%, due to low abundance and technical factors (46, 47). Additionally, we observed that the number of cells for each cell type is extremely imbalanced in dataset S0. For example, it contains 49,201 cancer cells but only 543 dendritic cells. To overcome these limitations and better mimic the bulk cell environment of solid tumors, we created dataset S1. We generated single-cell-type GEPs (scGEP) in dataset S1 by randomly sampling 100 cells from a cluster of cells with a specific cell type in dataset S0 and aveaging their expression values in transcript per million (TPM) format, normalized to sum to 1 million for each GEP in non-logarithmic space. Each generated GEP still has 1 million total expression values. We repeated this step for each cell type, resulting in dataset S1 (see **Table S1**) with 10,000 scGEPs for each of the 11 cell types.

### Pre-processing of the TCGA dataset

Since most of the single cell RNA-seq datasets were collected from patients with carcinoma, we selected 17 bulk RNA-seq datasets of carcinoma from TCGA. In addition, two brain tumors, Low-grade glioma (LGG) and Glioblastoma multiforme (GBM), were selected as the non-carcinoma controls (negative controls). For the 19 types of cancer selected in this study, CPE values were available for each tumor sample except Pancreatic adenocarcinoma (PAAD) (27). These values were used as approximations of the cancer cell proportions in each tumor sample. We converted raw read counts of downloaded GEPs to TPM format and only kept protein-coding genes for further analysis. We obtained the length and type of each gene from the annotation file “gencode.gene.info.v22.tsv” which was used in the TCGA project. The 19 types of cancer considered in this study were as follows: ACC, BLCA, BRCA, GBM, HNSC, LGG, LIHC, LUAD, PAAD, PRAD, CESC, COAD, KICH, KIRC, KIRP, LUSC, READ, THCA, and UCEC. The full names of these cancer types can be found from https://gdc.cancer.gov/resources-tcga-users/tcga-code-tables/tcga-study-abbreviations.

### Synthetic tumors with known cell proportions

To generate the bulk GEP of each synthetic tumor, we first need to define a cell proportion matrix (**P ϵ** ℝ^*m×n*^), where each column of the matrix corresponds to the proportion of one cell type (11 cell types in total), *m* denotes the number of synthetic tumors to be simulated, and *n* denotes the number of cell types considered.

### Random sampling

For each synthetic tumor *g* (one row in the cell proportion matrix) and each cell type *c* (one column in the cell proportion matrix), where *g ϵ {*1, …, *m}* and *c ϵ{* 1, …, *n }*, we randomly and independently sampled the cell proportion 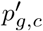from a uniform distribution *U* (0, 1), and then normalized the sum of each row to 1 using:

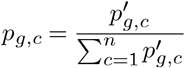

The result is a cell proportion matrix **P** with *m* rows and *n* columns.

### Segment sampling

For each synthetic tumor *g* and each cell type *c* included in the simulation, where *g ϵ{*1, …, *m}* and *c ϵ{*1, …, *n }*, we sampled the first cell proportion *p*_*g*,1_ from a uniform distribution *U* (0, 1). Then, for the following cell proportions *c ϵ {*2, 3, …, *n −* 1*}*, we sampled *p*_*g,c*_ from a uniform distribution 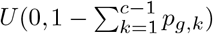, as long as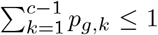. Once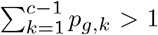, we assigned 0 to all the following cell types. Finally, we set the last column of each row to be 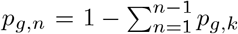 to ensure that 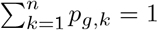 After repeating these steps *m* times, we obtained a cell proportion matrix **P** with *m* rows and *n* columns. To remove the bias caused by sampling order, we randomly shuffled the *n* values in each row of **P**.

### Gene expression profiles of synthetic tumors

In order to train our DNN model, we generated 3 datasets, which we refer to D0, D1, and D2. Each dataset contains *m* rows (bulk GEPs of synthetic tumors) and *q* columns (genes). These datasets were used as the training sets in **Fig. 2**. To ensure consistency with the integrated scRNA-seq dataset and the TCGA data, we used around 12,000 genes common to both datasets in the initial steps for generating the GEPs of synthetic tumors. For more details on each dataset, please refer to **Table S1**.

To generate **Di** (*i ϵ*{0, 1, 2}), we first generated a matrix of cell proportions, represented as **P**^**i**^ with *m*rows and *n* columns, where *m′* is the number of tumors to be synthesized and *n* is the number of cell types considered. Note that *m′* should be equal to *m* if there is no GEP-level filtering step, which will be further explained in the following part.

To generate a bulk GEP for one synthetic tumor based on single-cell-type GEPs in dataset S1, we compute the weighted sum of the GEPs of *n* cell types in TPM format, where the weights for each cell type are the corresponding cell proportions stored in each row of the matrix **P**^**i**^.

More specifically, for each row *g* in dataset *Di* representing a synthetic tumor *g*, we calculate:

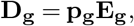

where **p**_**g**_ is a row vector with *n* elements (corresponding to a row in **P**^**i**^), and **E**_**g**_ is an *n* by *r* matrix (each row denotes a GEP for a specific cell type, and each column denotes a gene). The cell proportions in **P**^**i**^ were generated using the sampling methods described in the previous section, and the order of cell types in the rows of **E**_**g**_ matches the order of cell types in the columns of **P**^**i**^. Each row of **E**_**g**_ is randomly sampled from dataset S1 according to the corresponding cell type.

### GEP-level filtering

GEPs can vary in a huge highdimensional space without any constraint, but the valid GEPs of solid tumors only occupy a relatively small area within that space. Therefore, when simulating the GEPs of bulk tumors based on the integrated scRNA-seq data, the high-density areas of GEPs between synthetic tumors and real tumors may not match (as shown in the last column of **Fig. 2a**). To obtain a high-quality synthetic dataset, it is necessary to apply a filtering step based on the distribution of GEPs in real tumor samples. More specifically, we first constructed a median GEP (mGEP) across all 19 tumour types in the TCGA dataset. The expression value of each gene in the mGEP equals the median of gene expression values across 7,699 samples. The similarity between each generated GEP and mGEP was measured by the *L*^1^ norm. Only if the 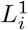 distance between the GEP *i* of a synthetic tumor and the mGEP is less than a specific threshold *r*(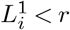), the GEP *i* will be included in the training set. To determine *r*, we calculated the *L*^1^ norm distance between mGEP and each GEP of the 7,699 samples, 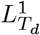, where *d ϵ {*1, 2, …, 7699*}. r* was set to the 95th quantile of 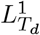 (Supplementary **Fig. S13**). Under such GEP-level filtering, on average, we can obtain 1 eligible GEP from every 20 generated GEPs.

To ensure that the final synthetic dataset contains enough high-quality GEPs, we repeated the cell proportion generation and GEP-level filtering steps multiple times. Specifically, we generated a cell proportion matrix with a certain number of rows, and then performed the GEP-level filtering to obtain a certain number of eligible GEPs. This process was repeated multiple times until we obtained the desired number of GEPs for the final synthetic dataset.

For example, to generate a dataset with 10,000 GEPs through GEP-level filtering, we can first generate a cell proportion matrix with 1000 rows and then perform the filtering to obtain around 50 eligible GEPs. By repeating these steps around 200 times to obtain 10,000 eligible GEPs in total. Note that the number of eligible GEPs obtained after each filtering step may vary due to the random sampling process and the threshold used for the *L*^1^ norm distance.

### Gene-level filtering

We also noticed that even after GEP-level filtering, the expression level of certain genes in the synthetic dataset can still be highly different from the corresponding gene expression level in the TCGA datasets. This problem may be due to the sparsity of scRNA-seq data. To further improve the similarity between synthetic dataset and TCGA, we performed gene-level filtering. There are two sequential steps for the gene-level filtering (using the synthetic dataset D1 as an example). Step 1): For each gene across all samples in D1, we calculated the Pearson correlation coefficient (*r*) between the expression values (TPM used) and the cell proportions of each cell type. Only those genes that their correlation coefficients were higher than 0.3 in at least one cell type were kept, and only the top 1000 most correlated genes were kept if more than 1000 genes existed for a specific cell type. We selected 0.3 as a threshold to ensure a proper number of genes were kept for each cell type. Step 2): For each gene, we compared the gene expression value range (log space) across all samples in D1 to that of the equivalent gene in the TCGA dataset. Only the genes that existed in both datasets were kept. We used the [*q*_5_, *q*_95_] quantile range of a gene *i* in the TCGA dataset, represented as 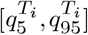, and the median expression level of the gene in D1 dataset, represented as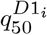, to determine if the gene should be kept.

Specifically, we kept the gene if and only if 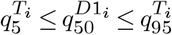, meaning that the expression level of the gene in the synthetic dataset has a large overlap (at least 55%) with the corresponding gene in the TCGA dataset.

After gene-level filtering, about 6000 genes were kept and then the GEP of each synthetic tumor was scaled to TPM.

As shown in the **Table S5**, the known marker genes for each cell type generally show the highest correlation with the corresponding cell type (highlighted by yellow, the marker genes of epithelial cells were used by cancer cells).

### DNN model. Model architecture

We used a fully connected multiple layer perceptron (MLP) with 5 hidden layers (Supplementary **Fig. S14**). Compared to the previous deep learning models, such as DigitalDLSorter (24) (2 layers) and Scaden (25) (4 layers), our model is deeper and narrower. This allows us to decrease the total number of parameters to prevent overfitting and expedite training while still being able to learn complex features. Notable, we intentionally used 100 units for the first hidden layer, which dramatically reduced the number of parameters without impairing the model’s performance.

### Model training and testing

Two datasets, D1 and D2, were combined to train DeSide (**Table S1**), which contains a total of 200,000 bulk GEPs (100,000 for each dataset). The common genes in both datasets were used (6,168 genes). All GEPs were converted to log_2_(*TPM* + 1) and then divided by 20 to ensure that all values were less than 1 before being used as input. Early stopping with “patience=100” was employed to prevent overfitting, and the maximum epoch was set to 3000 (usually, the training process stopped within 1000 epochs). The weighted average of MAE (mean absolute error) and RMSE (root-mean-square error) was used as the loss function, i.e., *αMAE* + (1−*α*)*RMSE*, where *α* = 0.8. The learning rate was set to 2*e* 5, and the batch size was set to 128. During training, we used 80% of the samples in the training set for model training, while the remaining 20% samples were used as a validation set. After training, the model’s performance was evaluated using 4 different test sets: T0 to T3 (see **Table S1**).

### GEP embeddings in the last (first) hidden layer of a DNN model

First, a DNN model *M* is trained by a specific training set. After training, the output layer (the layer following the last hidden layer) of the DNN model is removed to create a new model *M′* The output of *M′* is the GEP embeddings of the last hidden layer, which are low-dimensional vectors. Similarly, we can remove all the layers after the first hidden layer to create a new model, and its output is the GEP embeddings in the first hidden layer.

### Algorithm comparison

We compared DeSide with several available algorithms: EPIC, CIBERSORTx (CIBERSORT), Scaden and Kassandra.

#### EPIC

We used the R package of EPIC (version 1.1.5) with default parameters to infer cell proportions for the 19 types of cancer considered in this study. In the default setting, Tref (Tumor infiltrating cells profile) was used as the reference dataset. All gene expression values were converted to TPM for input.

### CIBERSORT/CIBERSROTx

We used the web-server of CIBERSORTx to infer cell proportions of 19 types of cancer. Two different single-cell datasets were used as reference datasets to create the signature matrix: HNSCC (provided by CIBERSROTx) and LUAD (the scRNA-seq dataset “luad_kim_05”, collected by this study). All gene expression values were converted to TPM for input. Since CIBER-SORTx uses the same algorithm as CIBERSORT to infer cell proportions of different cell types, we do not distinguish strictly between the two.

#### Scaden

We used the Python package of Scaden (version 1.1.2). Two different training sets were used to train the Scaden model: the ascites dataset (the only cancer-related dataset provided by Scaden) and simu_bulk (the combined dataset D1+D2 in this study). We followed the online document of Scaden to implement all steps (process, train and predict).

Note that the same dataset, simu_bulk, was used to train the DeSide model.

#### Kassandra

We used the web-server of Kassandra (solid tumor model) to predict cell proportions of all related cancer types. All gene expression values were converted to TPM for input.

#### Comparing with GSE184398 (Fig. 3c)

We scaled the provided flow scores from 0 to 100 to the range [0, 1] by dividing by 100 (the maximum flow score). The cell proportions of each cell type were scaled by dividing the maximum proportion of the corresponding cell type. The cell proportions of T cells were compared with the total cell proportions of CD4^+^ T cells and CD8^+^ T cells. Myeloid cells were compared with the total proportions of DC, macrophages, neutrophils, and mast cells if they were present. Stromal cells were compared with the total proportions of fibroblasts and endothelial cells if they were present. All null values were removed during the analysis. We excluded 4 cancer types, namely GBM (Glioblastoma), PNET (Primitive neuro-ectodermal tumors), SARC (Sarcoma), and MEL (Melanoma), that do not originate from epithelial cells, and we also excluded 6 samples with flow scores greater than 50 (6/74) from the analysis of stromal cells. The scRNA-seq dataset originated from HN-SCC (provided by CIBERSORT) was used to generate the signature matrix used in CIBERSORT. The ascites dataset (provided by Scaden) was used to train the Scaden model.

We used TensorFlow (48) (version 2.8.0) to construct and train DeSide. Single cell dataset processing was carried out using Scanpy (49) (version 1.8.0) and batch-effects were removed using BBKNN (50) (version 1.5.1). To facilitate the analysis, the expression values were transformed into logarithmic scale (log_2_(TPM+1)), and stored in the h5ad file format by anndata (version 0.8.0, available at https://github.com/scverse/anndata). The Wilcoxon signedrank test was performed by Scipy (51).

### Evaluation metrics. Signature score

For each cell type, the signature score was calculated by averaging the expression values of corresponding marker genes (**Table S4**). All GEPs were normalized to TPM before calculating the average values.

We used the following metrics to compare two vectors *x* and *y*, each of which has *N* elements.

### The Pearson correlation

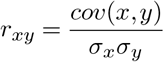

where the covariance is computed as 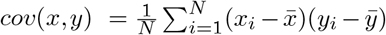, and the mean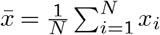, and the standard deviation 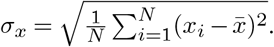.

### Concordance correlation coefficient (CCC) (52)

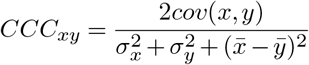

the same notations were used as in the Pearson correlation.

### Root mean square error (RMSE)

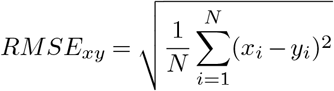

For one output of DeSide *y* = [*y*_1_, …, *y*_*k*_], where *k* = 10, the number of total cell types without cancer cell.

### Sigmoid function

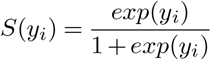

where *i ϵ{*1, …, 10}. Each element can be calculated independently.

### Softmax function

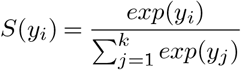

where *i ϵ {*1, …, 11} and *k* = 11. The sum of *y* is always 1. The Pearson correlation with p-value and RMSE was calculated by the Python library, SciPy (51) and scikit-learn (53) respectively.

### Survival analysis

We used the cell proportions predicted by DeSide to group the patient samples in 10 tumor types (COAD, KIRP, CESC, BLCA, LIHC, HNSC, KICH, LUAD, LUSC, READ) in the TCGA dataset, and then examined their prognosis. Each tumor type was selected by the following criterion: the correlation between predicted cancer cell proportions and corresponding CPE values (27) is higher than 0.6 and the prediction error, RMSE, is lower than 0.15. More specifically, to consider the impact of only one cell type on the patients, we first separated the patients into two groups (low and high) using the median value of predicted cell proportions as the threshold. To consider the two cell types together in the patients, we divided the patient samples into four groups (low-low, low-high, high-low and high-high) based on the median values of their proportions. If two of these four groups can be clearly identified as the best- and worst-survival groups, and the log-rank test p-value of these two groups is less than 0.05, we considered these two cell types as potential candidates for an interaction pair that may help us further screen patients’ prognosis in the clinical setting.

The survival information for individual patients were all obtained from TCGA (https://portal.gdc.cancer.gov/repository). Before we performed the statistical analysis, patients whose “days to death” and “days to last follow up” were both smaller than 1 day were removed from further analyses. The survival analysis was performed by Kaplan-Meier survival estimators, which is provided by the Python library lifelines (version 0.26.4). Any of two patient groups were compared by log-rank tests, and multiple groups (*>* 2) were tested by multivariate log-rank test. In this work, all the Hazard Ratio (HR) show in the figure were calculated by dividing the hazard rate of better survival group by the hazard rate of worse survival group. So in here, HR less than 1 suggests a lower risk in better survival group compared to worse survival group.

## ACKNOWLEDGEMENTS

We thank Dr. LAW Ellie Yuen Yi and Dr. Adam George CRAIG for critical feedback and discussion on the manuscript. This work was supported by the National Key R&D Program of China (2021YFA0911100), the National Natural Science Foundation of China (32170672 and 32000886), and the Guangdong Basic and Applied Basic Research Foundation (2021A1515012461).

## Data availability

All datasets used in this study are publicly available. The bulk RNA-seq data in GEO184398-Live (29) was downloaded from https://ftp.ncbi.nlm.nih.gov/geo/series/GSE184nnn/GSE184398/suppl/GSE184398_pancan_all_pc_genes_Live_TPM_Aug_3_20.tsv.gz. The Archetype Flow Populations values (i.e., flow scores) used in **Fig. 3c** were downloaded from https://datalibrary.ucsf.edu/node/121/.

We downloaded the bulk RNA-seq data of 19 tumor types in the TCGA project from GDC Data Portal (https://portal.gdc.cancer.gov/) as read counts (ht-seq.counts). The annotation file “gencode.gene.info.v22.tsv”, which contains the information of gene length, was downloaded from https://gdc.cancer.gov/about-data/gdc-data-processing/gdc-reference-files.

## Code availability

DeSide is implemented in Python using the TensorFlow library to build the DNN model. The source code is available on GitHub at: https://github.com/OnlyBelter/DeSide.

## Supplementary Note 1: Supplementary Information

**Fig. S1.**
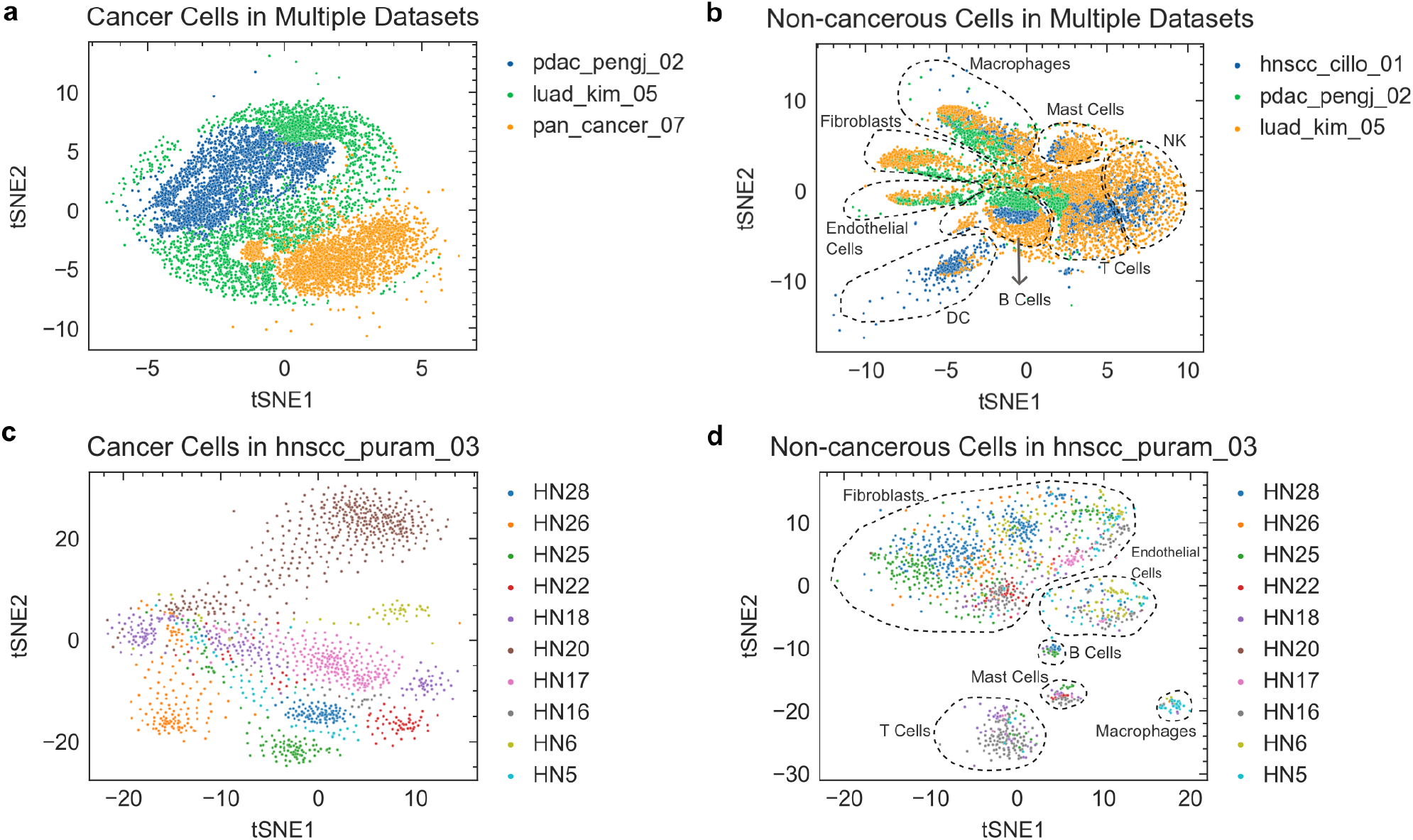
Comparison of gene expression profile variance between cancer cells and non-cancerous cells across different datasets or patients. **a**, t-SNE visualization of cancer cells in 3 different datasets from the integrated scRNA-seq dataset S0 (Supplemental **Table S2**). Each dataset has a small overlap with the other datasets. **b**, t-SNE visualization of non-cancerous cells in 3 different datasets in SO. Different datasets often have a big overlap for most of the non-cancerous cell types, which indicates less variance across different datasets compared to cancer cells. **c**, t-SNE visualization of cancer cells in the HNSCC dataset (Supplemental **Table S2**). Each patient tends to separate from each other. **d**, t-SNE visualization of non-cancerous cells in the HNSCC dataset. These non-cancerous cells that originated from different patients but with the same cell type tend to mix together, which indicates less variability across patients compared to cancer cells. For (c) and (d), the same patients were shown as in **Fig. 2a** and **2c** of (Puram et al., 2017) (39).

**Fig. S2.**
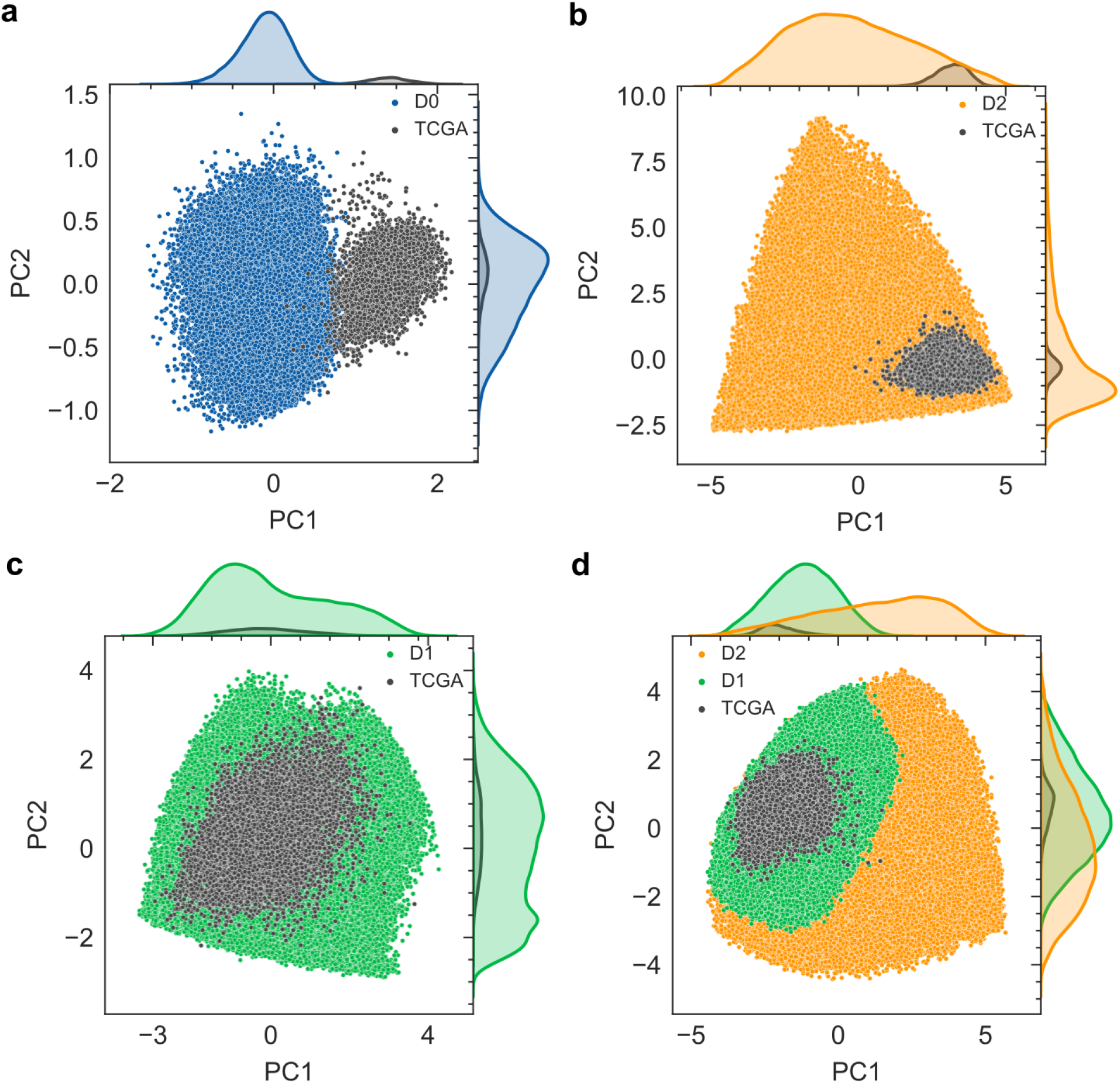
PCA visualization of the output of the first hidden layer in the DNN model. **a**, Model trained on dataset D0, which is separate from TCGA. **b**, Model trained on dataset D2. Despite D2 covers the entire area taken by TCGA, the high-density zones are separate. **c**, Model trained on dataset D1. Compared to b, D1 and TCGA show higher similarity after filtering. **d**, Model trained on D1 and D2.

**Fig. S3.**
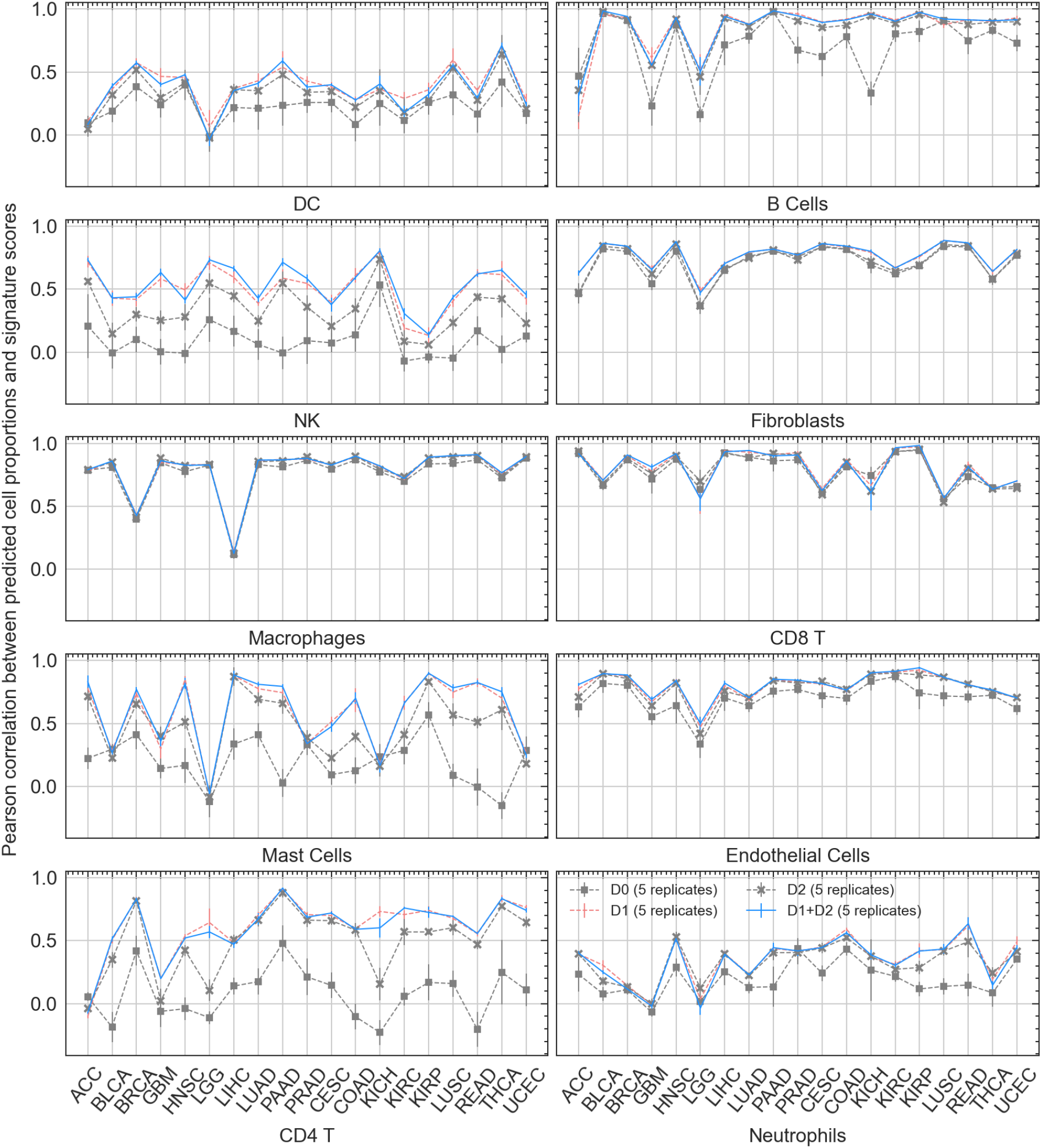
Pearson correlation between predicted cell proportions and the corresponding signature scores for each cell type across 19 cancer types. The signature score of each cell type was calculated by averaging the expression values of marker genes (**Table S4**). DNN model was trained by different training sets, D0, D1, D2 or D1+D2.

**Fig. S4.**
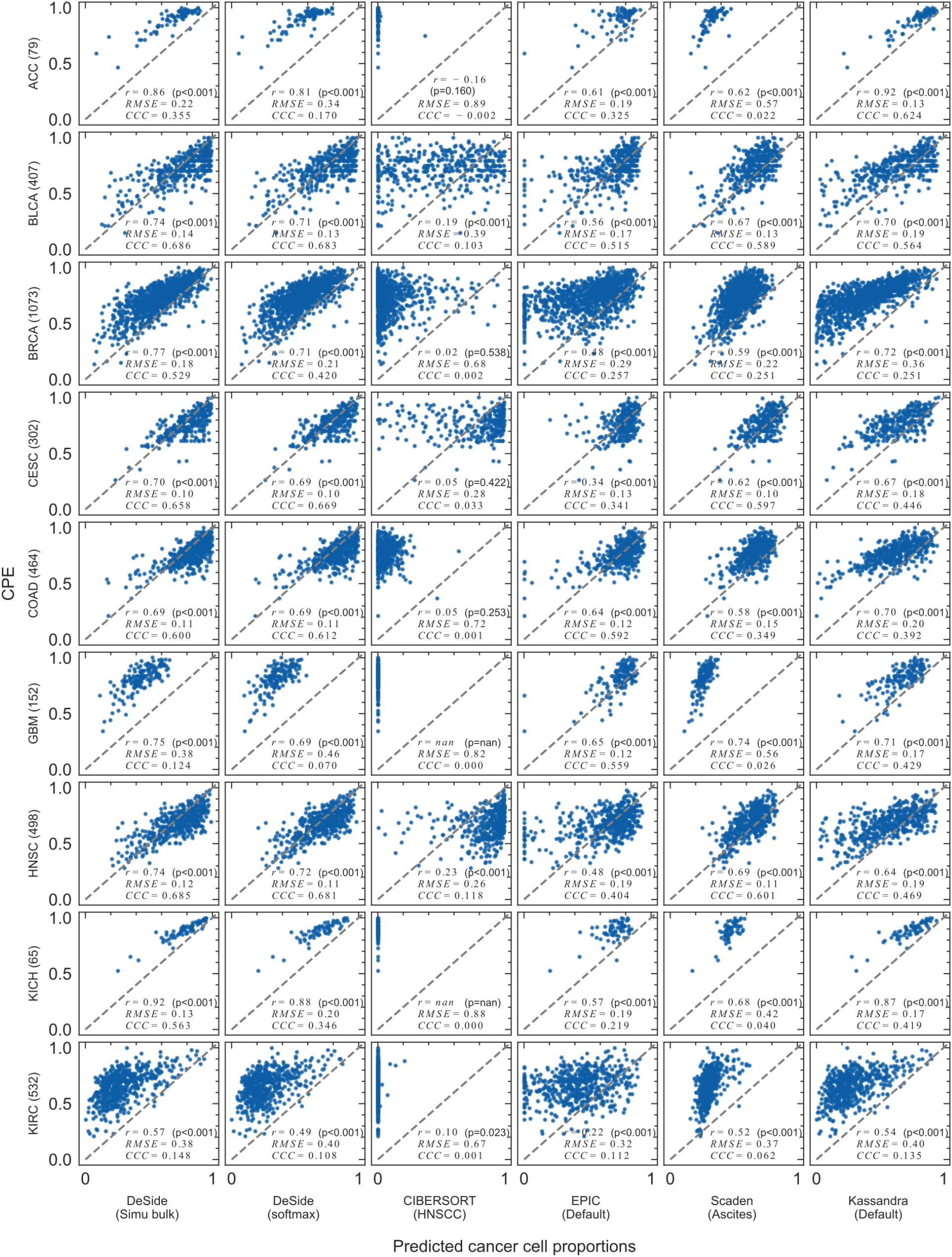
Comparison of predicted cancer cell proportions and the corresponding CPE values across 18 cancer types, part 1. 9 cancer types, ACC, BLCA, BRCA, CESC, COAD, GBM, HNSC, KICH, and KIRC, were compared. The cell proportions were predicted by 7 different algorithms (or datasets). The number of samples for each cancer type was shown in the parentheses of y-axis, and the dataset used for each algorithm was shown in the parentheses of x-axis. The Pearson correlation (*r*), root-mean-square error (RMSE), and concordance correlation coefficient (CCC) were shown on each subplot for a specific algorithm and a specific cancer type.

**Fig. S5.**
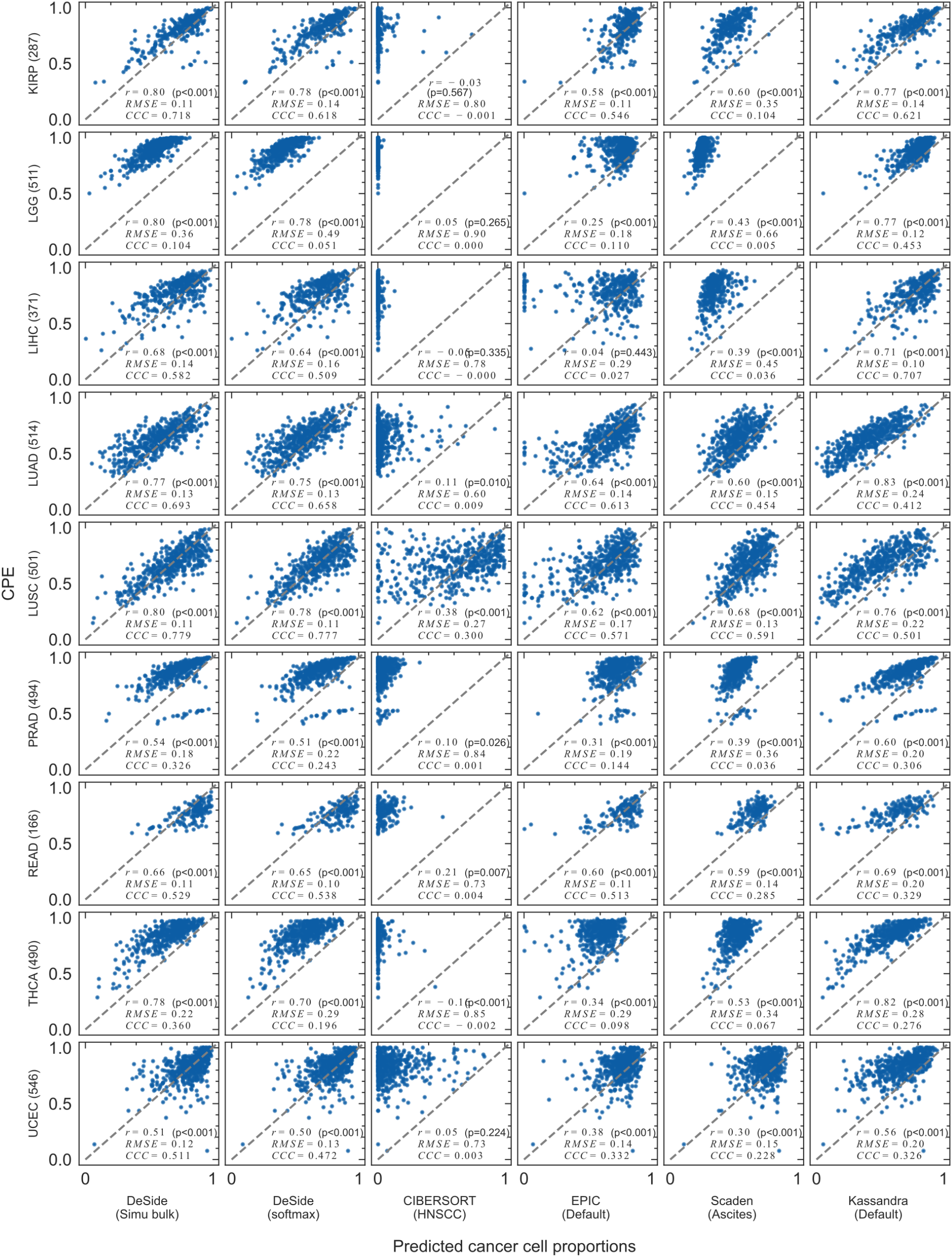
Comparison of predicted cancer cell proportions and the corresponding CPE values across 18 cancer types, part 2. 9 cancer types, KIRP, LGG, LIHC, LUAD, LUSC, LRAD, READ, THCA, and UCEC were compared. The cell proportions were predicted by 7 different algorithms (or datasets). The number of samples for each cancer type was shown in the parentheses of y-axis, and the dataset used for each algorithm was shown in the parentheses of x-axis. The Pearson correlation (*r*), root-mean-square error (RMSE), and concordance correlation coefficient (CCC) were shown on each subplot for a specific algorithm and a specific cancer type.

**Fig. S6.**
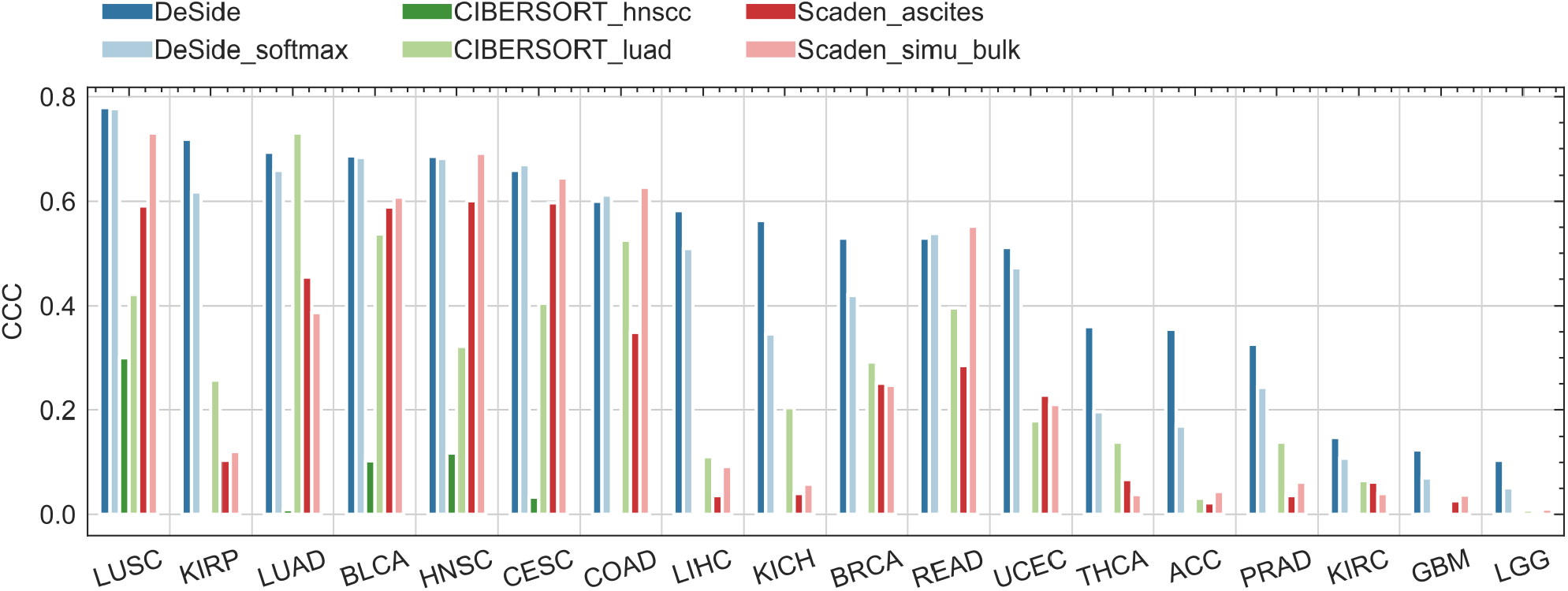
Comparison of concordance correlation coefficients (CCCs) across different activation functions or datasets. The effects of two different activation functions, sigmoid and softmax, on the output layer of DeSide’s DNN architecture were compared (dark blue and light blue bars). Additionally, the effects of different datasets were compared for both CIBERSORT (deep green and light green bars) and Scaden (deep red and light red bars). The suffix “_softmax” indicates that the softmax function was used as the activation function of the output layer in the DNN model. The suffix “_simu_bulk” indicates that the dataset D1+D2 was used as the training set. The suffix “_ascites” indicates that the ascites dataset provided by Scaden was used as the training set. The suffixes “_hnscc” or “_luad” indicate that the scRNA-seq data originated from HNSCC (provided by CIBERSORT) or LUAD (provided by this study) was used as the reference dataset to generate the signature matrix. Further details on the algorithm comparison can be found in the Methods section.

**Fig. S7.**
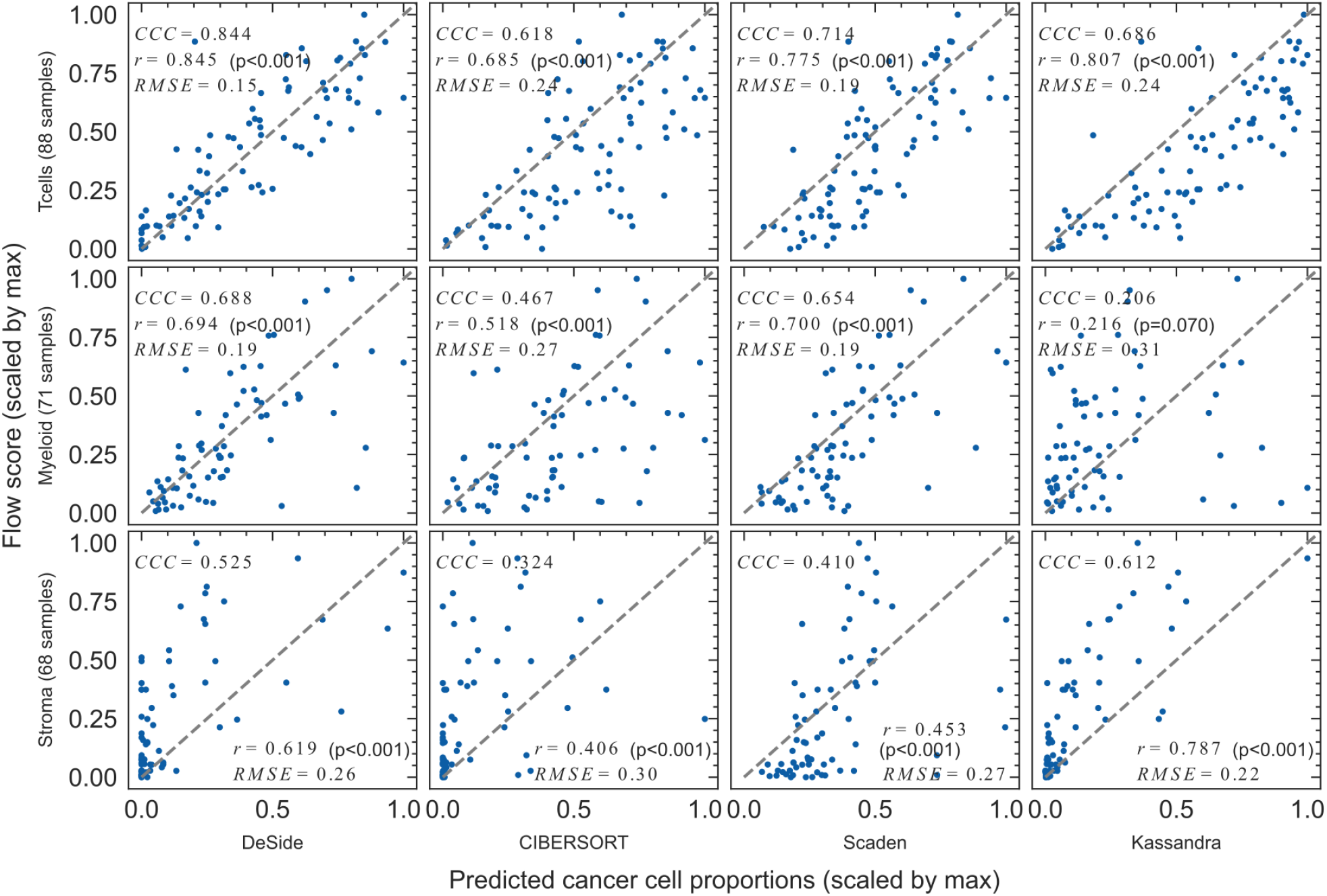
The scatter plot of Fig. 3c. The cell proportions of 3 cell types were predicted by 4 different algorithms and compared with flow score (measured by experiments). Each row shows one cell type: T cells, myeloid cells and stromal cells. All null values of flow scores were removed and the number of samples left for each cell type was shown in brackets. The 3rd row shows the result of stromal cells after removing 6 outliers. Both the predicted cell proportions and flow scores for each cell type were normalized by dividing the corresponding maximum values (see Methods).

**Fig. S8.**
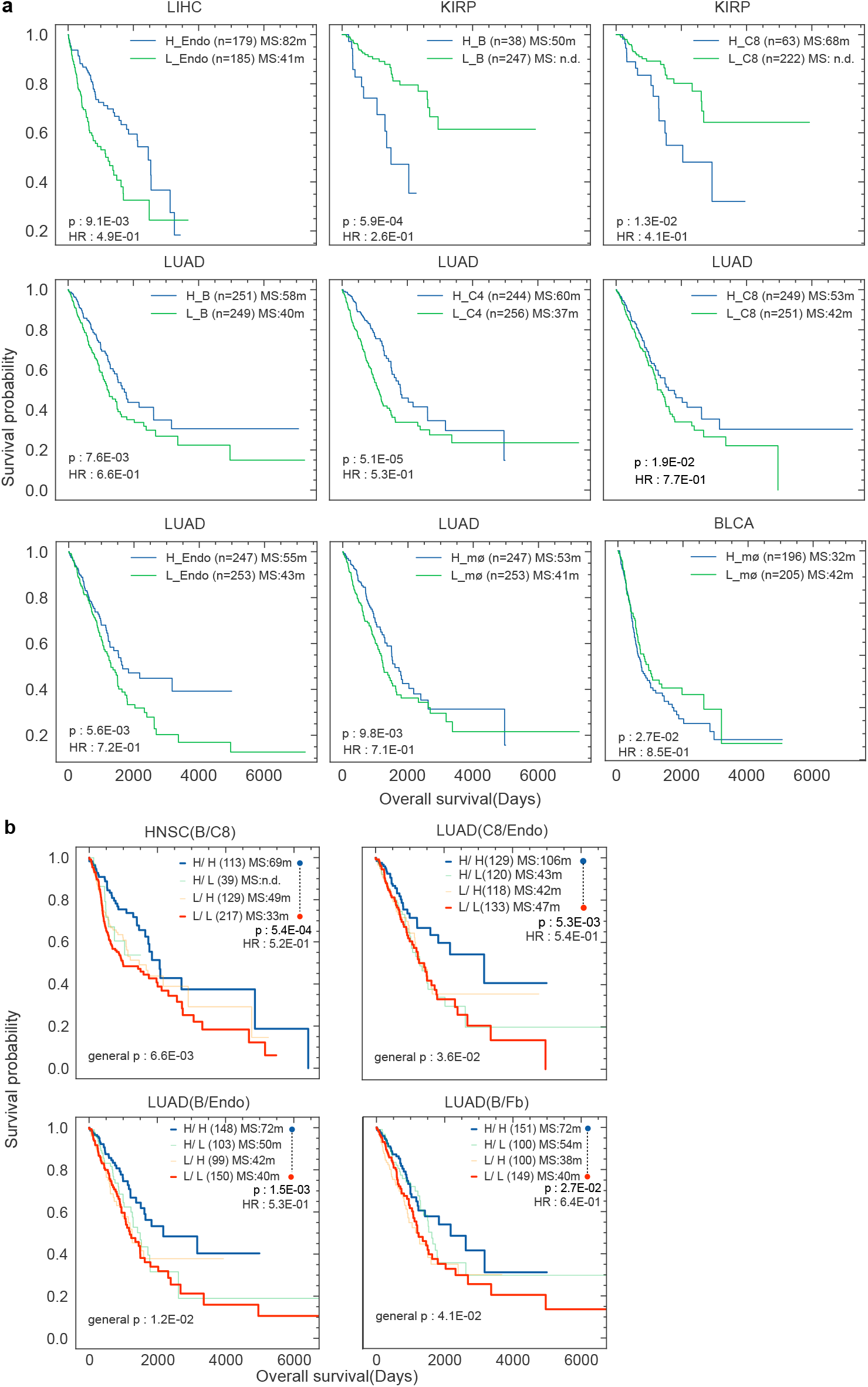
Kaplan-Meier overall survival (OS) curves for TCGA cohort classified by the cell proportions predicted by DeSide. **a**, OS was significantly different when classified by the cell proportions of a single cell type, which is not show in **Fig. 4a**.The p-values were calculated by log-rank test. **b**, OS was significantly different when classified by the cell proportions of two cell types, which is not show in **Fig. 4b**. One of the cell types in each pair can also separate patients into two groups and was shown in (a) or (**Fig. 4a**). The bottom left p-values were used to test the overall difference of the four survival curves, while the top right p-values were used to test the difference between the best and the worst survival curves. The log-rank test was used for all statistical significance testing. The numbers in parentheses are sample numbers. MS, Median survival; measured in months (m); n.d., not determined; HR, Hazard Ratio; Endo, Endothelial cells; C8, CD8^+^ T cells; mø, Macrophages; Fb, Fibroblasts; C4, CD4^+^ T cells; B, B Cells; H, High; L, Low.

**Fig. S9.**
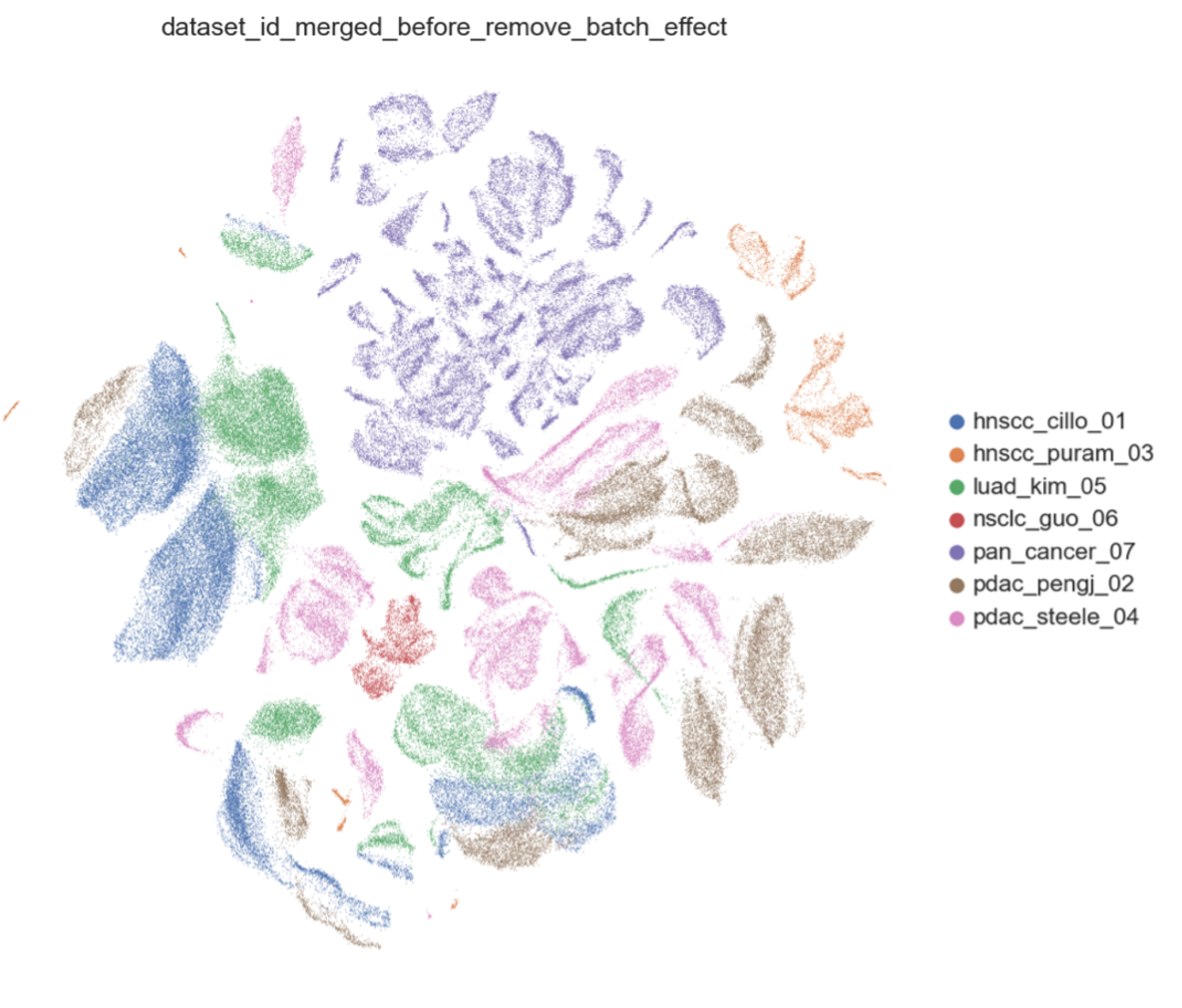
UMAP of 7 scRNA-seq datasets before removing batch effect across datasets. The cells were colored by the corresponding dataset. Since the existence of batch effect across different datasets, the cells were clustered by the belonging dataset instead of cell type.

**Fig. S10.**
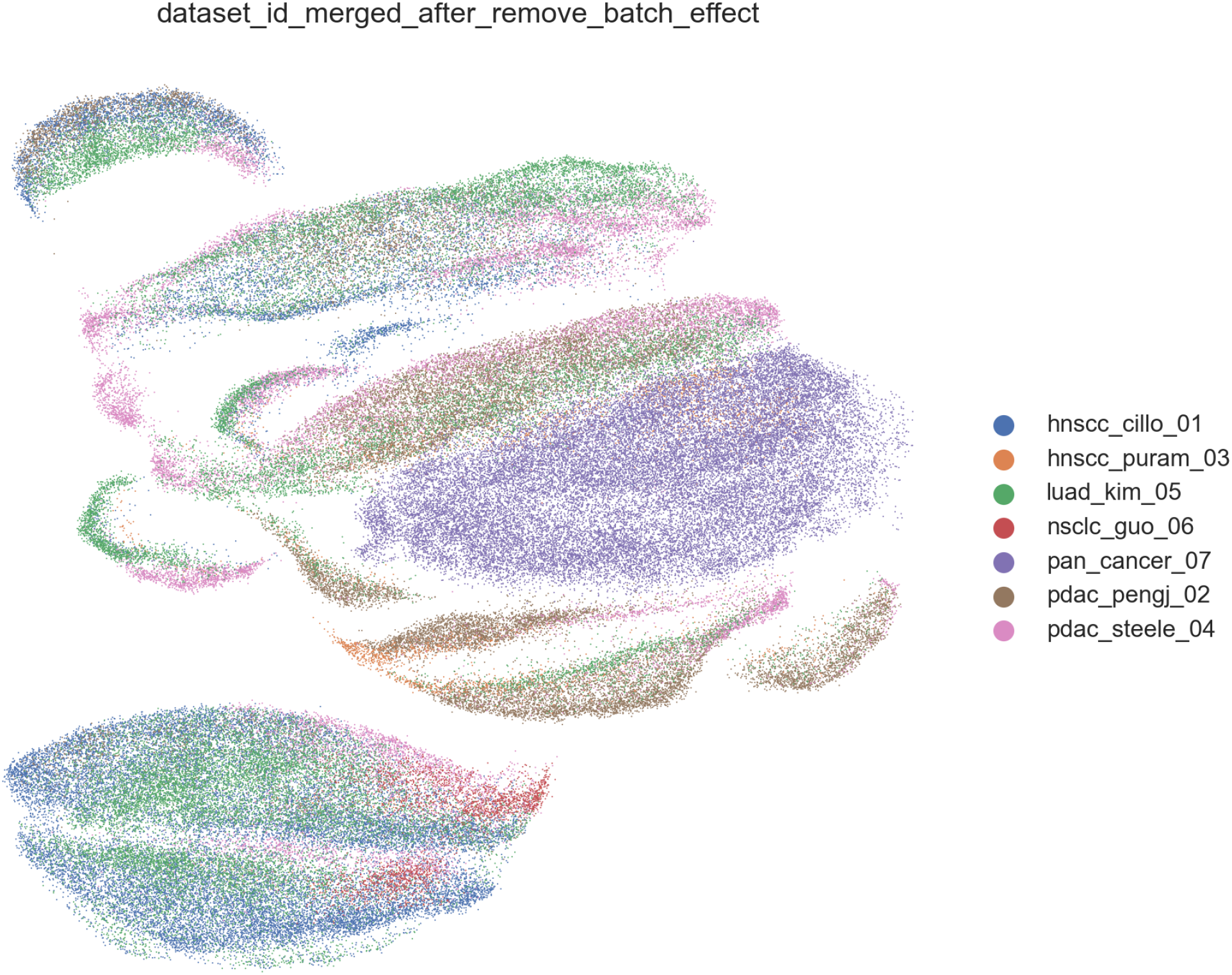
UMAP of 7 scRNA-seq datasets after removing batch effect across datasets. The cells were colored by the corresponding dataset. After removing the batch effect across different datasets, the cells that belong to the same cell type were clustered together. (Fig. S11)

**Fig. S11.**
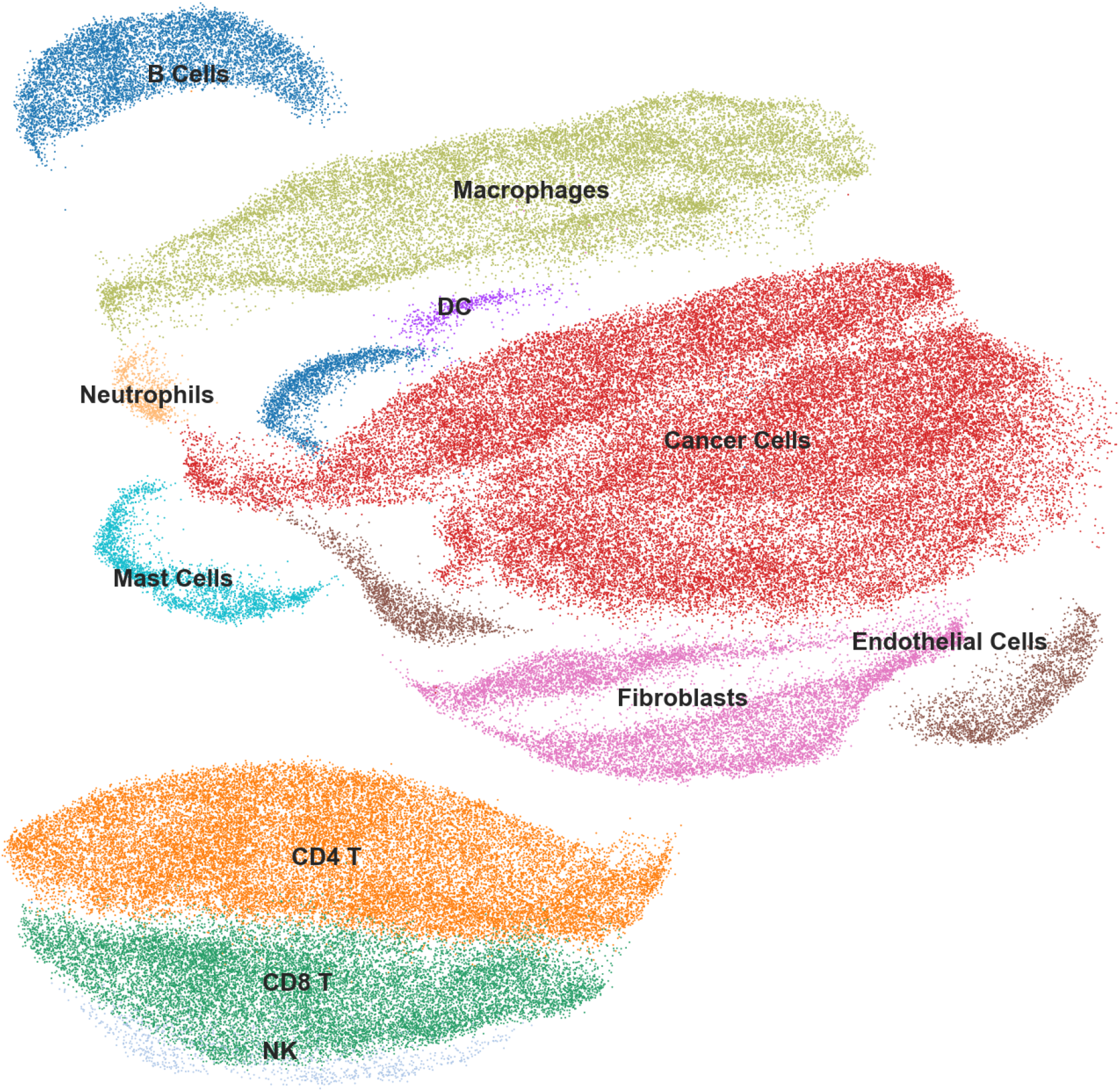
UMAP of 7 scRNA-seq datasets after removing batch effect across datasets. The cells were colored by the annotated cell type. The relative distance of different clusters could be meaningful to indicate the similarity between different cell types.

**Fig. S12.**
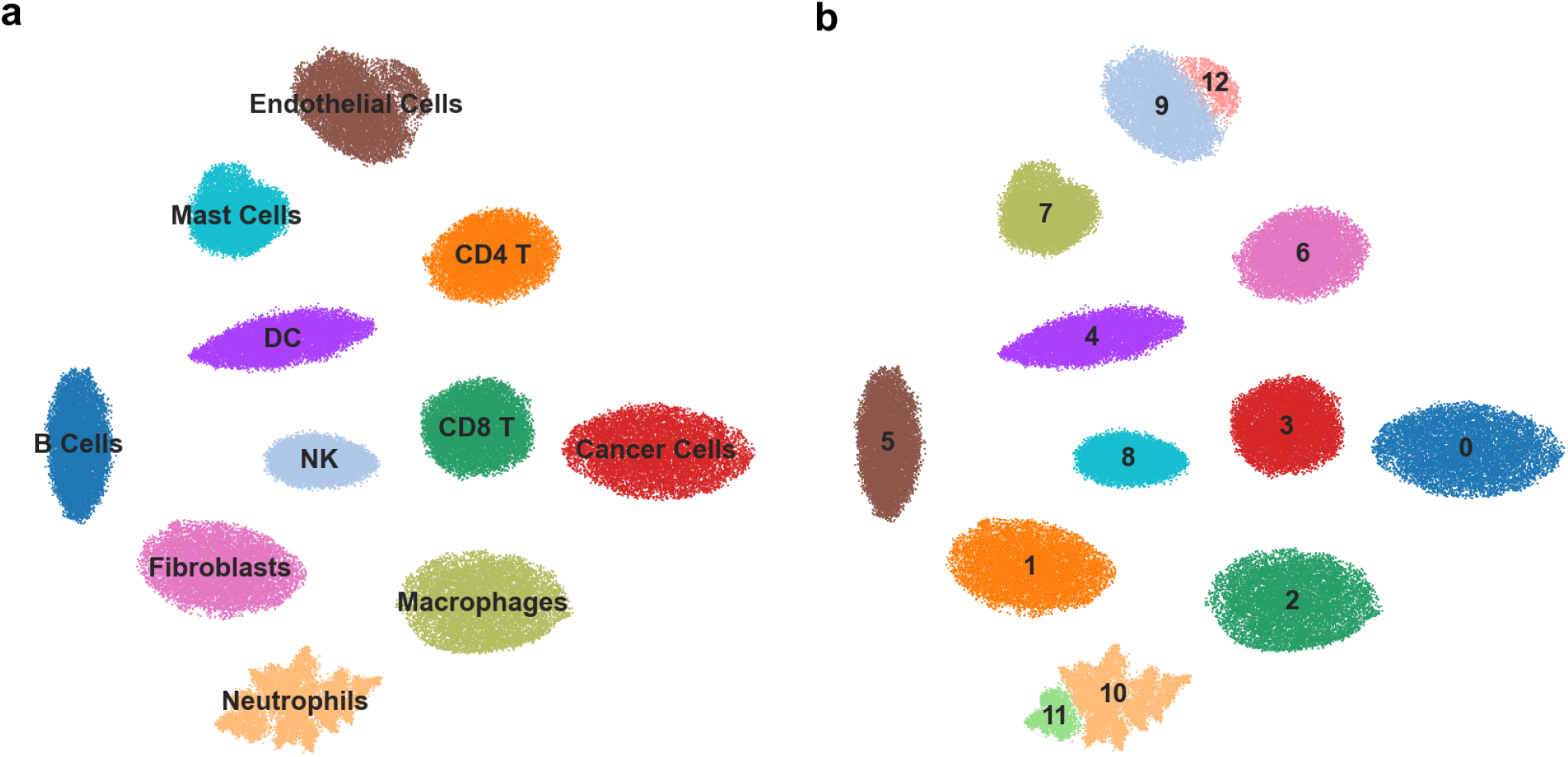
The UMAP visualization of dataset S1. **a**, UMAP was coloured by the cell type where each GEP was sampled from dataset S0. **b**, UMAP was coloured by cluster ids using Leiden clustering (54) (integrated in Scanpy (49)). Neutrophils and endothelial cells were clustered into 2 different but adjacent sub-clusters respectively.

**Fig. S13.**
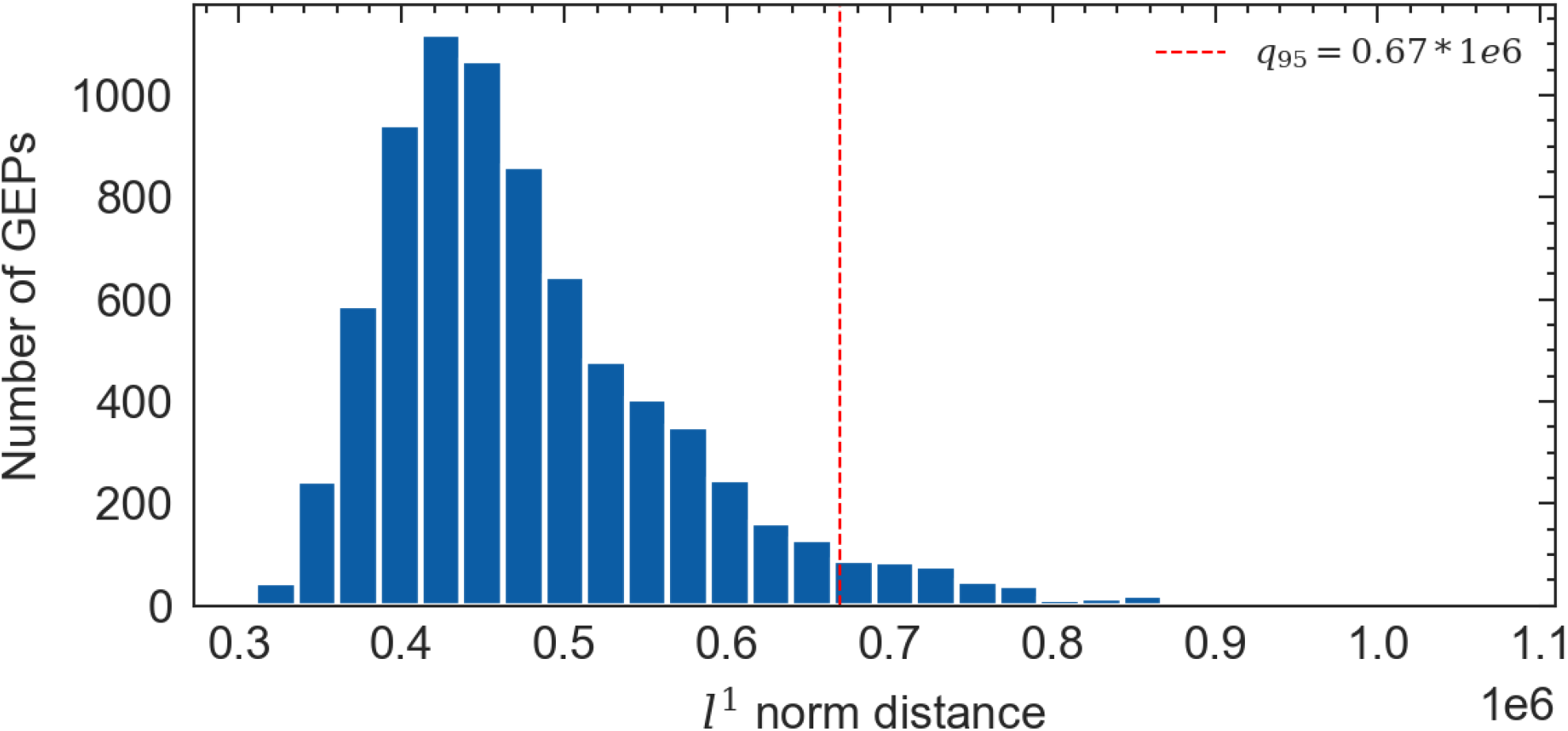
The histogram of *l*^1^ norm distance between median GEP (mGEP) and 7,699 samples in TCGA dataset, 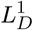. The red vertical line represents the 95% quantile of 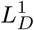.

**Fig. S14.**
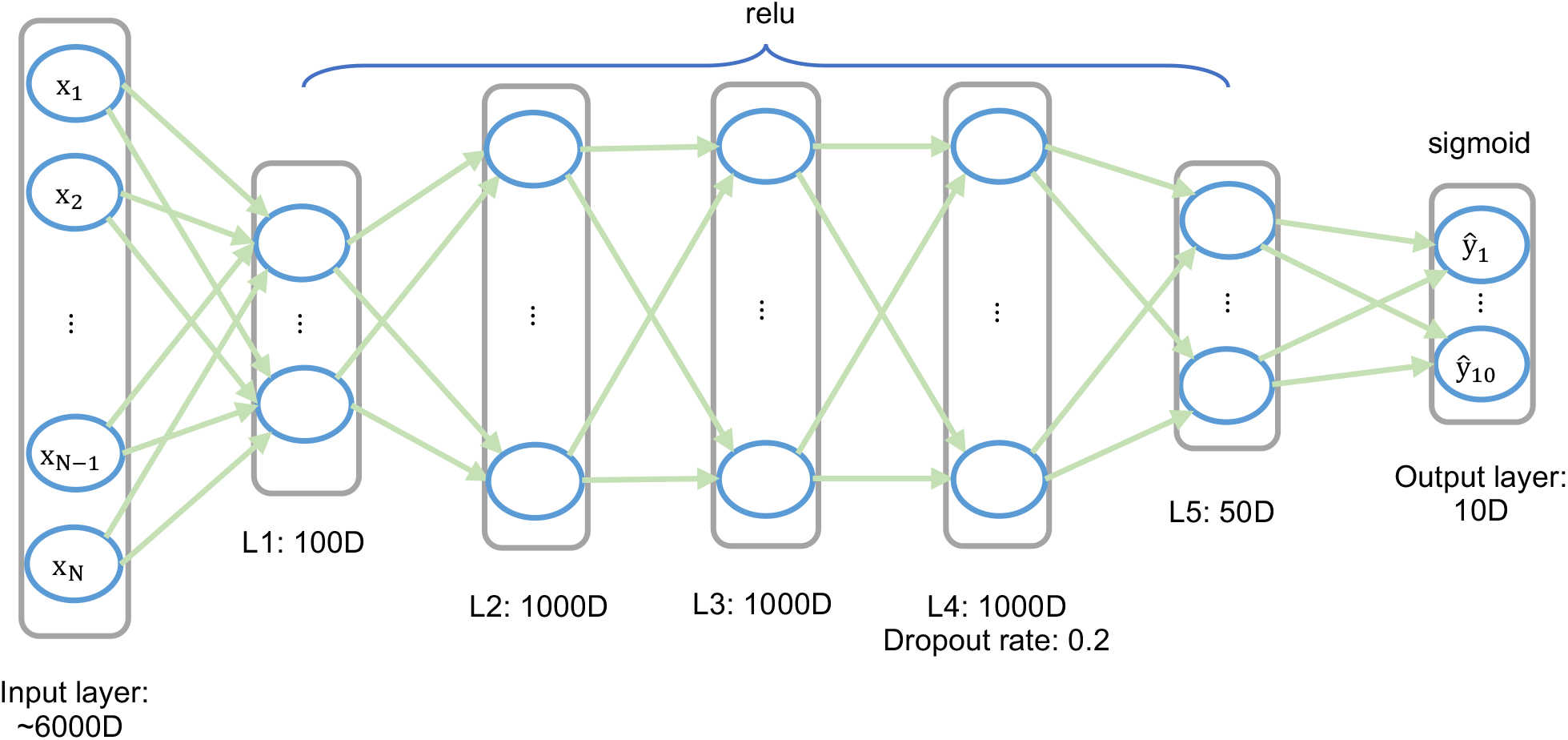
The architecture of DNN model in DeSide. The gene expression profiles with about 6000 genes were imported by input layer. Relu activation function was used for all hidden layers and sigmoid activation function was used in the output layer. A dropout layer was added after layer 4 with 0.2 dropout rate.

